# Deciphering spatio-temporal fluorescence changes using multi-threshold event detection (MTED)

**DOI:** 10.1101/2020.12.06.413492

**Authors:** Franziska E. Müller, Volodymyr Cherkas, Gebhard Stopper, Laura C. Caudal, Laura Stopper, Frank Kirchhoff, Christian Henneberger, Evgeni G. Ponimaskin, Andre Zeug

## Abstract

Recent achievements in indicator optimization and imaging techniques promote the exploration of Ca^2+^ activity patterns as a main second messenger in many organs. Astrocytes are important regulators of brain activity and well known for their Ca^2+^-dependent modulation of neurons. However, standardized methods to analyze and interpret Ca^2+^ activity recordings are missing and hindering global comparisons. Here, we present a biophysics-based concept to analyze Ca^2+^signals, which includes multiple thresholds and provides the experimenter with a comprehensive toolbox for a differentiated and in-depth characterization of Ca^2+^ signals. We analyzed various *ex vivo* and *in vivo* imaging datasets and verify the validity of our multi-threshold event detection (MTED) algorithm across Ca^2+^ indicators, imaging setups, and model systems from primary cell culture to awake, head-fixed mice. Applying our MTED concept enables standardized analysis and advances research using optical readouts of cellular activity to decrypt brain function. It allowed us to obtain new insights into the complex dependence of Ca^2+^activity patterns on temperature and neuronal activity.

**Highlights:**

→ We present a robust pixel-based algorithm to analyze multidimensional fluorescence data.
→ Automated multiple-threshold analysis accurately quantifies changes in fluorescence across magnitudes.
→ It characterizes the complexity of dynamic and overlapping activity patterns of Ca^2+^ activity of astrocytes *in vitro, in situ*, and *in vivo*.
→ Its application provides quantitative parameters how temperature and neuronal activity determine astrocytic Ca^2+^ activity.

## Introduction

Fluorescence microscopy is a method used across many scientific disciplines. In biology, it allows researchers to visualize cellular processes in time and space largely preserving the integrity of the sample. Recent developments in novel microscopy techniques, biosensors, and indicators push functional imaging to its limits. A persistent major challenge is to adequately analyze multidimensional data. For example, Ca^2+^ is an important signaling molecule and, because of its large concentration gradients, a well-established readout for cellular activity. In some cell types, such as astrocytes, Ca^2+^ activity patterns can be very complex. They are therefore a useful testbed for the development of analytical tools. Astrocytes are a crucial element in the brain architecture and display very distinct morphological characteristics that vary between brain regions and sub-regional networks^1^. With their fine processes astrocytes ensheathe synapses, providing structural support. As active partners at the tripartite synapse, they release gliotransmitters that regulate synaptic activity^2–5^. In return, the release of a variety of neurotransmitters has been shown to modulate the cytosolic Ca^2+^ concentration ([Ca^2+^]) in astrocytes^6^. Variations in astrocyte [Ca^2+^] have been proposed to represent a unique manner of cellular signaling, but the exact functions and regulatory mechanisms are only slowly emerging ^7,8^. Ca^2+^ signals in astrocytes are not solely stimulus-dependent, but also occur spontaneously^9^. They can also propagate and fuse, thus creating complex spatiotemporal patterns of activity. Whether these diverse patterns in Ca^2+^activity correlate to distinct astrocytic functions and morphological features is a matter of extensive debate.

Ca^2+^ activity in astrocytes appears as local and very distinct fluctuations, but could also spread into distant regions or engage the whole cell. Therefore, the spatial extent and amplitude of Ca^2+^ events are additional measures and standard evaluation approaches used in other applications cannot simply be transferred to astrocytic Ca^2+^ activity^10^.

A standard approach to analyzing Ca^2+^ activity is fluorescence microscopy using Ca^2+^-sensitive fluorescent dyes^11^. They generally come in two flavors: organic dyes, such as Oregon Green 488 BAPTA-1 (OGB-1), and genetically encoded Ca^2+^ indicators (GECIs), such as GCaMPs, which both have their specific advantages and disadvantages^8^. Both are easy to use in single channel recordings, which simplifies the readout and subsequent

data processing^12,13^. Various tools are available to extract relative changes in [Ca^2+^], which mainly analyze the frequency and magnitude of events^14^ based on predefined or automatically detected active regions^15,16^. Recently, alternative approaches have been presented that also analyze the directionality of propagating events and the origin of Ca^2+^ events, which is predominantly located in the fine peripheral processes^17,18^. This has shifted the focus from major Ca^2+^ changes in the somatic region to the periphery. However, these regions provide only a weak fluorescent signal due to their low volume. Therefore, when the detection method is based on identifying signals above noise level, events in these areas can escape detection due to a low signal-to-noise-ratio.

We developed a strategy to determine the signal of Ca^2+^ indicators at a basal [Ca^2+^], F_0_, which was the basis for calculating (F-F_0_)/F_0_, expressed as ΔF/F_0_, as a measure of relative changes in [Ca^2+^] in a pixel-based manner. We then applied an automated multi-threshold event detection (MTED) algorithm to quantify the intrinsic Ca^2+^activity of astrocytes in primary hippocampal cultures, organotypic slice cultures, and cortical astrocytes *in vivo*. Next, we tested MTED in two straightforward experimental settings. We could demonstrate substantially different Ca^2+^ activity dynamics of cultured astrocytes at room temperature (RT) compared to 37°C, which resembled the activity measured *in vivo*. We could further reveal that neuronal activity favors the generation of long-lasting [Ca^2+^] elevations in astrocytes. Our flexible MTED algorithm, exploited here successfully for Ca^2+^ indicators and astrocyte physiology across preparations in 2D over time, is easily applicable to investigate functional dynamics using various fluorescence-based indicators and biosensors and can be easily extended to 3D over time and to other cell types.

## Results

### Visualization of Ca^2+^ transients in astrocytes

For developing a strategy to accurately characterize Ca^2+^ activity, we used primary cultures of mouse hippocampal astrocytes expressing the Ca^2+^ indicator GCaMP6s. These cells exhibit extensive endogenous Ca^2+^ activity with varying amplitudes and magnitudes, changing directionality and regional patterns (Figure 1, Supplementary Movie 1). The GCaMP6s fluorescence signal F is sensitive to changes in Ca^2+^ levels, but also depends on the indicator concentration itself and the cellular structure, which determines the number of molecules in the focal volume. Figure 1 shows an example of Ca^2+^ changes in cultured hippocampal astrocytes. From the GCaMP6s fluorescence signal F (Figure 1a), which scales with both the Ca^2+^and GCaMP concentration, it is not possible to deduce the changes in [Ca^2+^] directly. The time series shown in Figure 1b contains various regions presenting similar brightness at some points over time (regions of interest (ROIs) 1-4). While the intensity of ROI 4 stays constant over time, F in ROIs 1-3 varies and can be identified as changes in Ca^2+^ levels. To obtain a quantity, which is independent from the indicator concentration and the number of indicator molecules in the focal volume, we developed an automated pixel-based algorithm for the determination of F_0_derived from the indicator signal F (Supplementary Figure 2 and Supplementary Movie 2). F_0_, deduced from the temporal behavior of F, represents the indicator signal at lowest [Ca^2+^], in the best case corresponding to darkest indicator brightness state possible. The resulting ΔF/F_0_provides an indicator concentration-independent readout which reflects relative changes in [Ca^2+^] (Figure 1c and e, Supplementary Movie 1). Alternatively, cytosolic co-expression of a Ca^2+^-insensitive fluorescent protein (FP; e.g., tdTomato) as a reference probe (F_R_) can be used to scale the indicator signal for pixel and time dependent differences in the number of indicator molecules in the focal volume (compare Supplementary Figure 1 and Supplementary Movie 1). A common strategy of detecting Ca^2+^ events from F is based on identifying signals above noise level (often a factor times the standard deviation of F). This can be a source of biases because only very pronounced changes in Ca^2+^ exceed the noise level at small, peripheral structures, where F is small as well (see Supplementary Figure 3). We can further visualize the spatiotemporal increases and decreases in cytosolic Ca^2+^ (*d*(ΔF/F_0_)/*d*t), depicted as red and blue regions, respectively, in Figure 1d and Supplementary Movie 1d.

**Figure 1:**
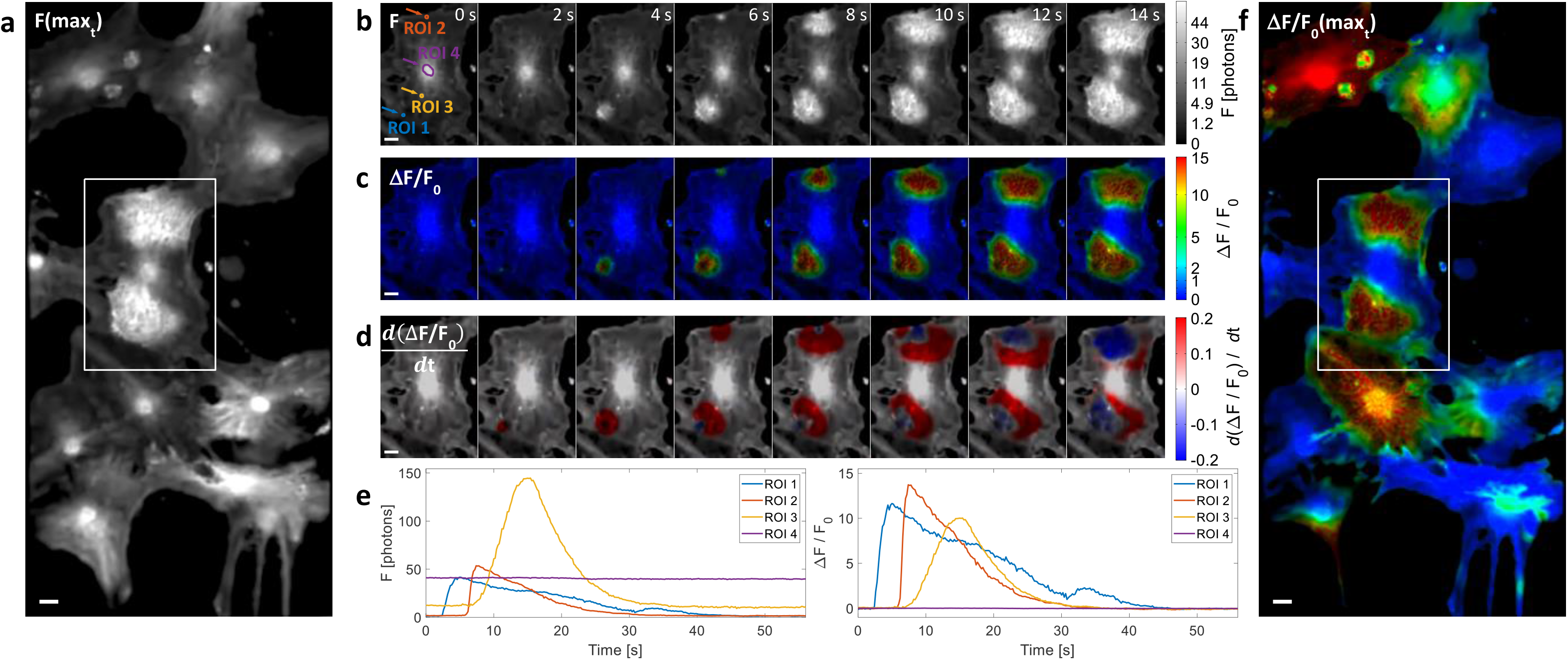
Visualization of Ca^2+^ activity using intensity-based indicators. **a**, Maximum fluorescence signal F (F(max_t_)) of Ca^2+^ indicator GCaMP6s in a 14 s sequence of a 10 min recording of primary mouse hippocampal astrocytes. Scale bar = 10 µm. **b**, Propagation of Ca^2+^ activity reflected by increased F intensity in the zoom region (white box in **a**). Various regions of interest (ROIs) show similar brightness at a given time point due to the GCaMP content in the confocal volume, but only those with changing Ca^2+^ levels vary in brightness over time (compare ROIs 1-3 with ROI 4). Scale bar = 10 µm. **c**, Application of ΔF/F_0_ by scaling F to lowest Ca^2+^ presence deduced from its temporal behavior to cancel out differences in F due to different number of indicator molecules in the focal volume. **d**, the color-coded amplitude of changes in Ca^2+^levels as *d*(ΔF/F_0_)/*d*t, with red colors indicating an increase in Ca^2+^ and blue colors representing decreasing Ca^2+^. **e**, Time trace of the selected ROIs shown in **b**, with dissimilar properties and different changes in F over time (compare **b-c**), obtained by the described approaches in **b-c. f**, Counterpart to **a** showing ΔF/F_0_(max_t_), which revealed profound spatial differences in detected Ca^2+^ activity.

Comparison of the overview images in Figure 1a and f, where the maximum projection of a 15 s time window is shown as intensity F and as false color ΔF/F_0_, respectively, underlines the benefit of the ratiometric concept. Bright regions, like some cell somata in Figure 1a, are characterized by low changes in Ca^2+^, whereas others exhibit high Ca^2+^ activity.

### Dynamic event detection

The analysis of Ca^2+^ activity in astrocytes should provide detailed information on event duration, size, and magnitude (see Figure 1, Supplementary Movie 1). A fundamental question is: which changes in Ca^2+^ can be called a Ca^2+^event? Furthermore, differences in the characteristics of Ca^2+^events need to be identified and should be reported by an event detection algorithm. Such an algorithm should also capture the spatial extent of the events without prior definition of static ROIs. Furthermore, the detection algorithm should not require experiment-specific settings. Therefore, we developed a pixel-based, dynamic event detection algorithm relying on multiple thresholds, which computes the abovementioned variables (Figure 2a and b, Supplementary Movie 3). Condition-specific adjustments of parameters are not required.

**Figure 2:**
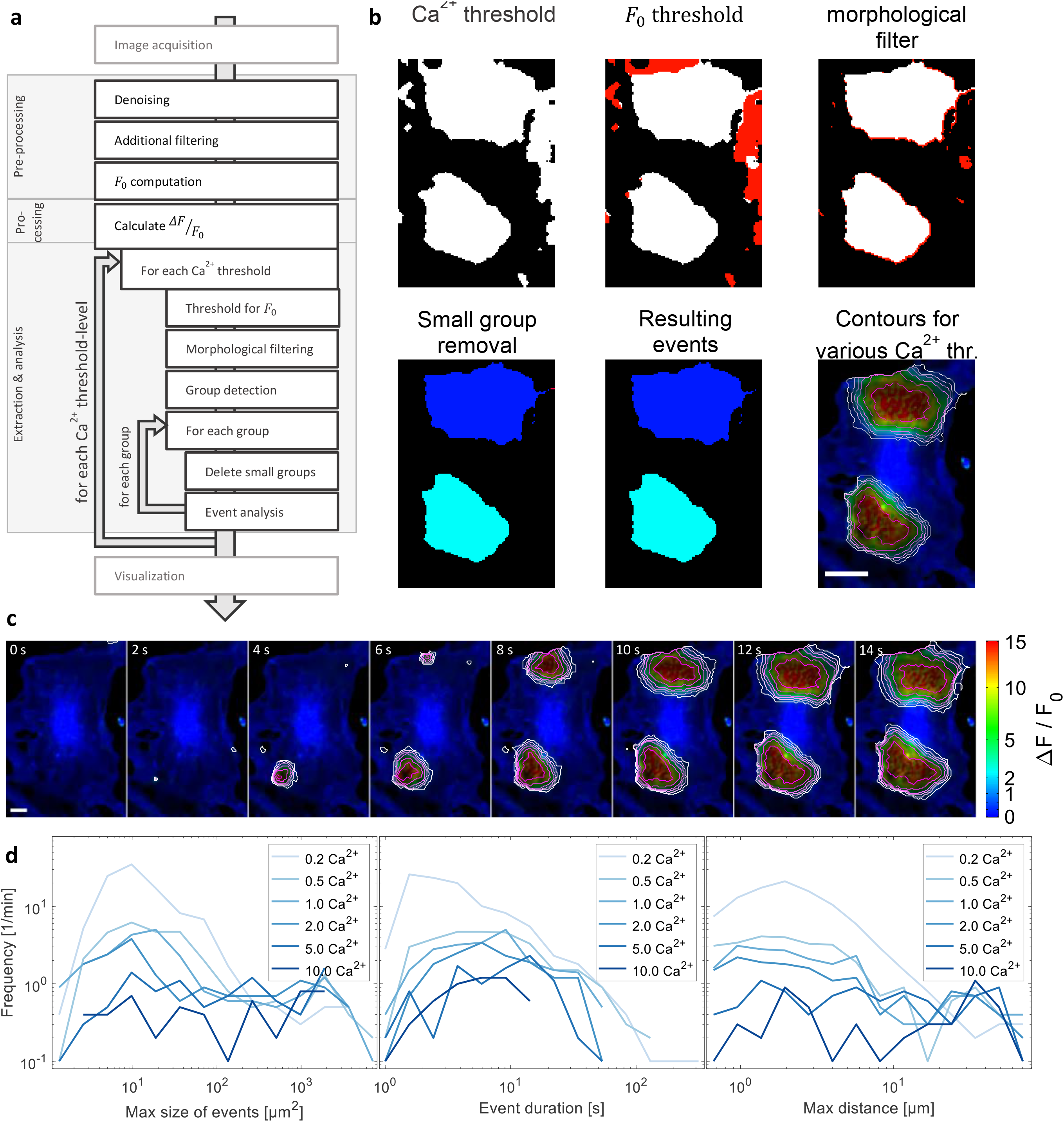
Data processing and Ca^2+^ event detection workflow. **a**, Diagram of steps in the evaluation process: image acquisition, data preprocessing, including a novel approach for a pixel-based F_0_ calculation, Ca^2+^ fluctuation recognition, ΔF/F_0_ and the multi-threshold approach for event detection (MTED), and the final activity visualization. **b**, Representative images of the main steps in the data processing workflow depicted in **a**. The red pixels depict regions removed from the previous step. Scale bar = 20 µm. **c**, Visualization of Ca^2+^ events calculated from the zoom region in Figure 1 and shown for various thresholds from Ca^2+^ = 0.2, 0.5, 1, 2, 5, 10 (white to magenta). Scale bar = 10 µm. **d**, Main output characteristics of the Ca^2+^ activity analysis: maximum size of detected events (lateral extent, x,y), mean event duration (t), and maximum distance the area center travelled within the event (µm).

Through Monte Carlo simulations, we found that the accuracy in identifying a pixel above a specific Ca^2+^ threshold can be far below 95% confidence and varies with F_0_(see Supplementary Figure 3). We selected a set of six thresholds (0.2, 0.5, 1, 2, 5, 10) expressed in terms of ΔF/F_0_, where a Ca^2+^ event reaching a threshold of 0.2 or 1 does not necessarily reflect a 20% or double increase in basal Ca^2+^ concentration, respectively (see Semyanov et al. 2020^8^). In conjunction with proximity relationships of neighboring pixels (xyt), MTED identifies groups of Ca^2+^-positive pixels for a set of Ca^2+^ thresholds, thereby allowing for dynamic region changes. The output for the sequence shown in Figure 1c is shown in Figure 2c and Supplementary Movie 4. Here, the six Ca^2+^ thresholds applied are depicted as contour lines with different colors, which show the respective spatial signal dynamics (Supplementary Movie 4c). A statistical analysis of the complete dataset is shown in Figure 2d as frequency plots for all analyzed Ca^2+^ thresholds exemplifying the maximum size of the events, their duration, and the maximum distance of the event center travelled over time (see Supplementary Table 1 for more statistical parameters). The frequency plots shown in Figure 2d can be accumulated for different experimental conditions and be compared as shown below. Overall, the MTED algorithm provides a comprehensive analysis of spatiotemporally complex fluorescence changes.

### The temperature-dependence of Ca^2+^ event characteristics

We next applied the analysis to two physiologically meaningful experimental scenarios. Investigations with cultured astrocytes are often carried out at RT (i.e., ∼ 25°C) rather than at the physiological body temperature (i.e., 37°C). When investigating Ca^2+^ event characteristics of the same astrocytic culture at different environmental temperatures, we observed substantial differences in the Ca^2+^ activity patterns (Figure 3, Supplementary Movie 5). Figure 3a illustrates the Ca^2+^ events detected within the zoomed-in region shown in Figure 1b-e for 25°C, 34°C, and 37°C. At 25°C, both events (ROI 1 and 2 from Figure 1) slowly increased, reaching large areas and high amplitudes. At 34°C, events in nearby regions were much shorter and occupied much smaller areas. At 37°C, the detected events that originated from the same initial region were spatially restricted and shorter. Such temperature-dependent changes become more obvious when illustrated in 3D (Figure 3b, Supplementary Movie 6). Statistical analysis of the maximum event size, event duration, and maximum distance of event propagation revealed that at 25°C strong Ca^2+^ changes up to 10-fold were detected, spanning over the complete parameter range for the maximal size and distance of Ca^2+^ events (Figure 3c, Supplementary Figure 4, Supplementary Movie 7). At 34°C, high-amplitude Ca^2+^ events were still detected but overall smaller amplitudes became more frequent. At 37°C, high Ca^2+^levels were not reached any longer and small-amplitude Ca^2+^events dominated. Event duration and maximum propagation distance also display a clear temperature dependence such that Ca^2+^ events become shorter and propagated less at higher temperature.

**Figure 3:**
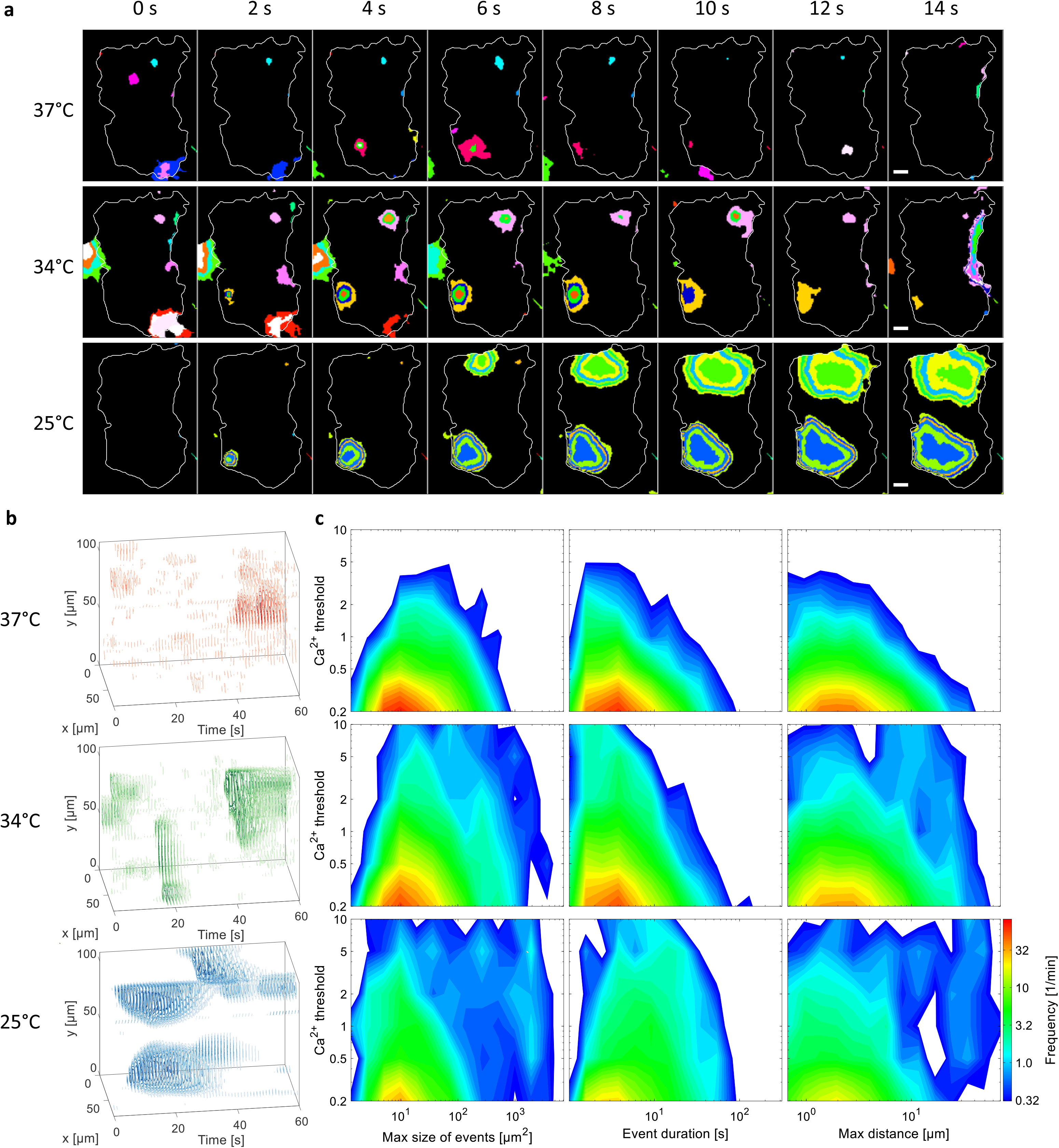
Astrocytic Ca^2+^ event characteristics are shaped by ambient temperature. **a**, Comparison of Ca^2+^ activity detected in the same cell at 37°C, 34°C, and 25°C for a sequence of 14 s color-coded for various thresholds. Data were captured in a time sequence of 10 min at 25°C, the sample heated to 34°C, and a sequence acquired from the same region. The measurement was repeated at 37°C and the sample cooled back to 34°C and then to 25°C (see Supplementary Movie 5). Scale bar = 20 µm. **b**, 3D contour plots of detected events, with darker colors representing higher changes in Ca^2+^, illustrating the change in Ca^2+^ activity with environmental temperatures. **c**, 2D histograms as a statistical presentation of selected parameters. At 37°C, the detected events are very small in space, transient in time, and do not reach high amplitudes. These characteristics change with cooler environments, reaching longer lasting, spatially more extensive, and comparably high amplitude events during measurement at 25°C.

### Suppression of cytosolic Ca^2+^ clearance reproduces the low temperature pattern

To understand the mechanisms underlying the observed temperature-dependent differences in the Ca^2+^ patterns, we compared the maximum increase and decrease in the Ca^2+^ signal, *max*_*t*_(*d*(Δ*F*/*F*_0_)/*dt*) and *min*_*t*_(*d*(Δ*F*/*F*_0_)/*dt*), respectively, for each Ca^2+^ event detected (Figure 4a). At 37°C, the centers of the signal increases were very local with low amplitude. In contrast, regions with positive changes in Ca^2+^ were spread over wider areas and reached greater amplitudes at lower temperatures. Similarly, the decrease in Ca^2+^ overlaying centers of maximal increase was punctual at 37°C but spread over wider regions at lower temperatures. Overall, negative Ca^2+^ changes were substantially greater at 34°C than 25°C. The rate of positive and negative Ca^2+^ changes was similar at 37°C, the positive Ca^2+^ changes (i.e., spreading of events) overcame the negative changes (i.e., depletion of events) at 25°C (Figure 4b).

**Figure 4:**
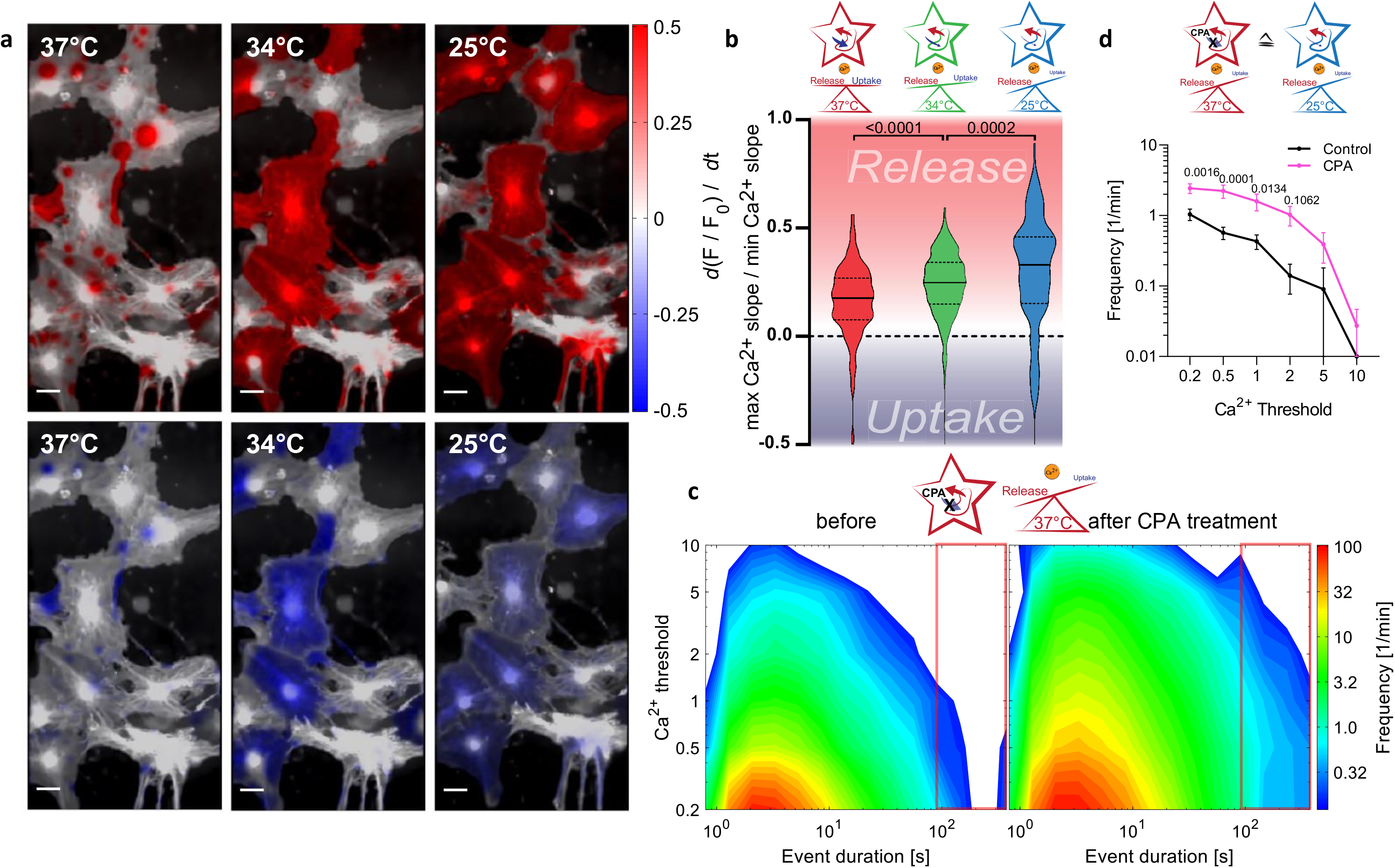
Slow physiological processes account for extended Ca^2+^ events at low temperatures. **a**, Visualized maximum increase *max*_*t*_(*d*(Δ*F*/*F*_0_)/*dt*) and decrease *min*_*t*_(*d*(Δ*F*/*F*_0_)/*dt*) of the Ca^2+^ signal reveals distinct behavior at the given temperatures. Scale bar = 20 µm. **b**, Ratio of the maximum and minimum Ca^2+^ change of all detected events in the measurements of the three investigated temperatures. For all Ca^2+^ thresholds, threshold 1 is shown, and the ratio reveals the discrepancy of Ca^2+^ release to its uptake velocity, which is increased at 25°C compared to 34°C and 37°C. **c**, 2D histograms of Ca^2+^ event duration at 37°C before and after incubation with Ca^2+^-ATPase inhibitor CPA. Pharmacological blockage of Ca^2+^ uptake led to similar event characteristics as obtained at low temperatures (n = 10, N = 3). **d**, Statistical evaluation of Ca^2+^ activity with duration >90 s as a function of the Ca^2+^ threshold applied in the algorithm. Data show mean and SEM. Two-way ANOVA with Sidak’s multiple comparisons post-hoc test.

Next, we used this analysis to investigate what underlies the profoundly different Ca^2+^ characteristics. The decrease in Ca^2+^ levels reflects its removal from the cytosol into stores of the endoplasmic reticulum (ER) and, thus, depends to a significant degree on Ca^2+^ pumps localized in the ER membrane (sarco/endoplasmic reticulum Ca^2+^ ATPases; SERCA pumps). Therefore, their inhibition should strongly affect Ca^2+^ activity patterns. Pharmacological inhibition of SERCA pumps by cyclopiazonic acid (CPA, 10 µM) strongly prolonged Ca^2+^ transients. The results obtained from these experiments suggest that the temperature-dependent slowdown of the ATP-driven Ca^2+^ uptake processes can explain the differences in the observed Ca^2+^ activity patterns (Figure 4c and d). Notably, lowering the temperature only partially slows down ATP-driven Ca^2+^uptake processes, whereas CPA blocks ATP-driven Ca^2+^ uptake to a greater extent. As in Figure 4b, analyses of the ratio of maximal positive to negative temporal Ca^2+^ changes were performed for each pixel of an event (Supplementary Figure 5). At low temperatures, the uptake is substantially blocked homogeneously throughout the cells. Overall, the MTED analysis presented here successfully captured complex signal patterns and enabled us to compare them between experiments.

### Event characteristics in situ

We next analyzed astrocytic Ca^2+^ events in organotypic slice cultures from the mouse hippocampus. In such preparations, the morphological organization of the hippocampus is mostly preserved, and the maturation of different cell types, network connections, and receptor/channel expression are closer to the situation *in vivo*^19,20^. Figure 5a and b show the average ΔF/F_0_ and the fraction of active time detected for the same preparation at 37°C and 25°C. Notably, in organotypic slices, we obtained a similar temperature-dependence of astrocyte Ca^2+^ characteristics as in primary astrocyte cultures. At 37°C, the maximal Ca^2+^ levels were lower and events shorter and spatially more restricted, whereas at 25°C, astrocytes exhibited long-lasting Ca^2+^events of lower frequency (Figure 5c). Detailed analysis of Ca^2+^event duration in primary astrocyte cultures and organotypic hippocampal slices at 37°C revealed an additional set of long-lasting high-magnitude events in the latter preparations (areas in open red boxes, compare Figure 5c to Figure 3c). As the main difference between these two preparations is the absence of neurons in primary astrocytic cultures, we hypothesized that long-lasting Ca^2+^ events may be a consequence of neuronal activity. Supporting this hypothesis, we observed a similar Ca^2+^ activity pattern in astrocyte-neuron co-cultures (Supplementary Figure 6). To verify the impact of neurons in the hippocampal slice preparations, we blocked neuronal activity by applying tetrodotoxin (TTX) (Figure 5d). This treatment led to a pronounced reduction in long-lasting Ca^2+^ events in both organotypic preparations and astrocyte-neuron co-cultures, shifting Ca^2+^ activity to an event pattern observed in primary hippocampal astrocyte cultures at 37°C (Figure 5e, Supplementary Figure 6). These findings demonstrate that the MTED algorithm is able to identify the neuronal impact on astrocytic Ca^2+^ activity.

**Figure 5:**
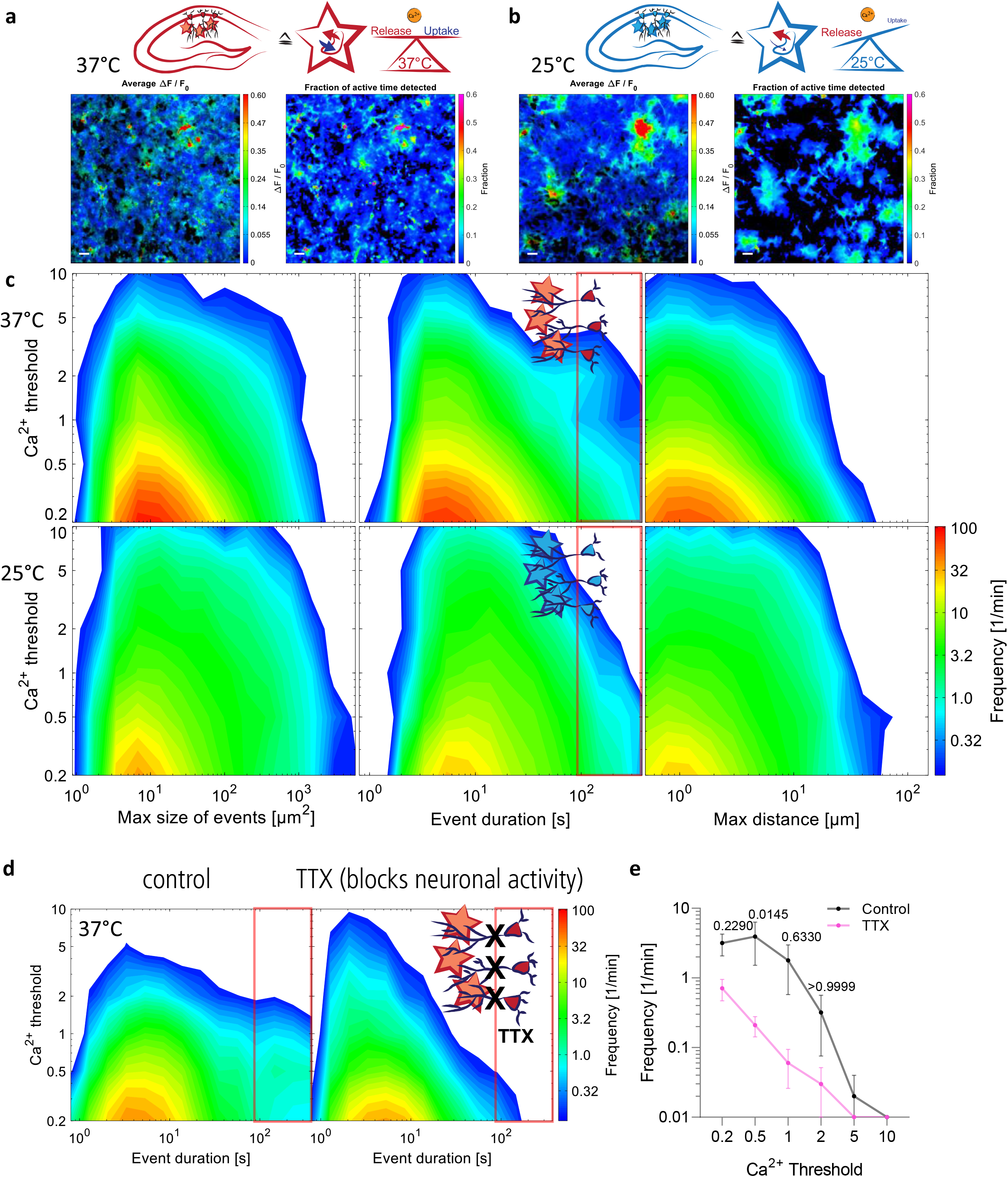
Temperature phenotype of Ca^2+^ activity is not restricted to primary astrocyte cultures. **a-b**, Average ΔF/F_0_ (left images) and the fraction of active time, which is the relative time for each pixel above a Ca^2+^ threshold of 0.2 (right images), detected in organotypic hippocampal slice cultures at 37°C (**a**) and 25°C (**b**). Comparably higher ΔF/F_0_ with spatially more extended and longer lasting events was detected at 25°C, similar to primary astrocyte cultures. Scale bars = 20 µm. **c**, 2D histograms of the selected parameters maximum size, duration, and travelled distance of events characteristic for organotypic slice cultures incubated at 37°C, and 25°C incubation temperature depict similar changes in features as observed in primary cultures. **d**, Averaged 2D histogram of control measurements in organotypic slice cultures showing prolonged event duration at 37°C compared to primary astrocyte cultures (red framed area, left panel). Blocking the neuronal activity with TTX reduces the occurrence of long-lasting Ca^2+^ events and exhibits similar 2D histogram features as primary astrocyte cultures at 37°C. **e**, Statistical evaluation of Ca^2+^ activity with duration > 90 s (red framed area in **d**) as a function of the Ca^2+^ threshold applied in the algorithm. Data show mean and SEM. Two-way ANOVA with Sidak’s multiple comparisons post-hoc test. n=10 recordings from 3 experiment days in **d** and **e**.

### Event detection in recordings obtained in vivo

To further investigate the versatility of the MTED approach, we evaluated astrocytic Ca^2+^ activity *in vivo*, for which the recorded fluorescence signal F is commonly much smaller, resulting in more challenging quantification of Ca^2+^ signals. To this end, we implanted a cortical cranial window into transgenic mice with astrocytic expression of GCaMP3. We set the excitation power between 30 – 40 mW to avoid light-induced Ca^2+^ activity and tissue damage, rarely detecting more than one photon per pixel and time point. Subsequently, we recorded and compared the Ca^2+^ activity patterns of the same cortical region in anesthetized and awake mice (Figure 6). Figure 6b depicts the average ΔF/F_0_ obtained, which coincides with the fraction of active time and shows overall low activity in the cortex of an anesthetized mouse. When the volatile anesthetic isoflurane was removed and mice woke up, the Ca^2+^ activity in the same region was drastically increased (Figure 6b, Supplementary Figure 7, Supplementary Movies 8 and 9). Our analysis revealed that, compared to anesthetized conditions, awake mice had higher magnitude Ca^2+^ events and increased frequency of long-lasting events (Figure 6c, Supplementary Figure 7c-e). Figure 6d displays the differences in the Ca^2+^activity pattern between anesthetized and awake conditions. Changes in frequency are most prominent at threshold levels 0.5 and 1 and for the subset of 1.5 - 7 s lasting events, which was supported by the statistical analysis (Figure 6e).

**Figure 6:**
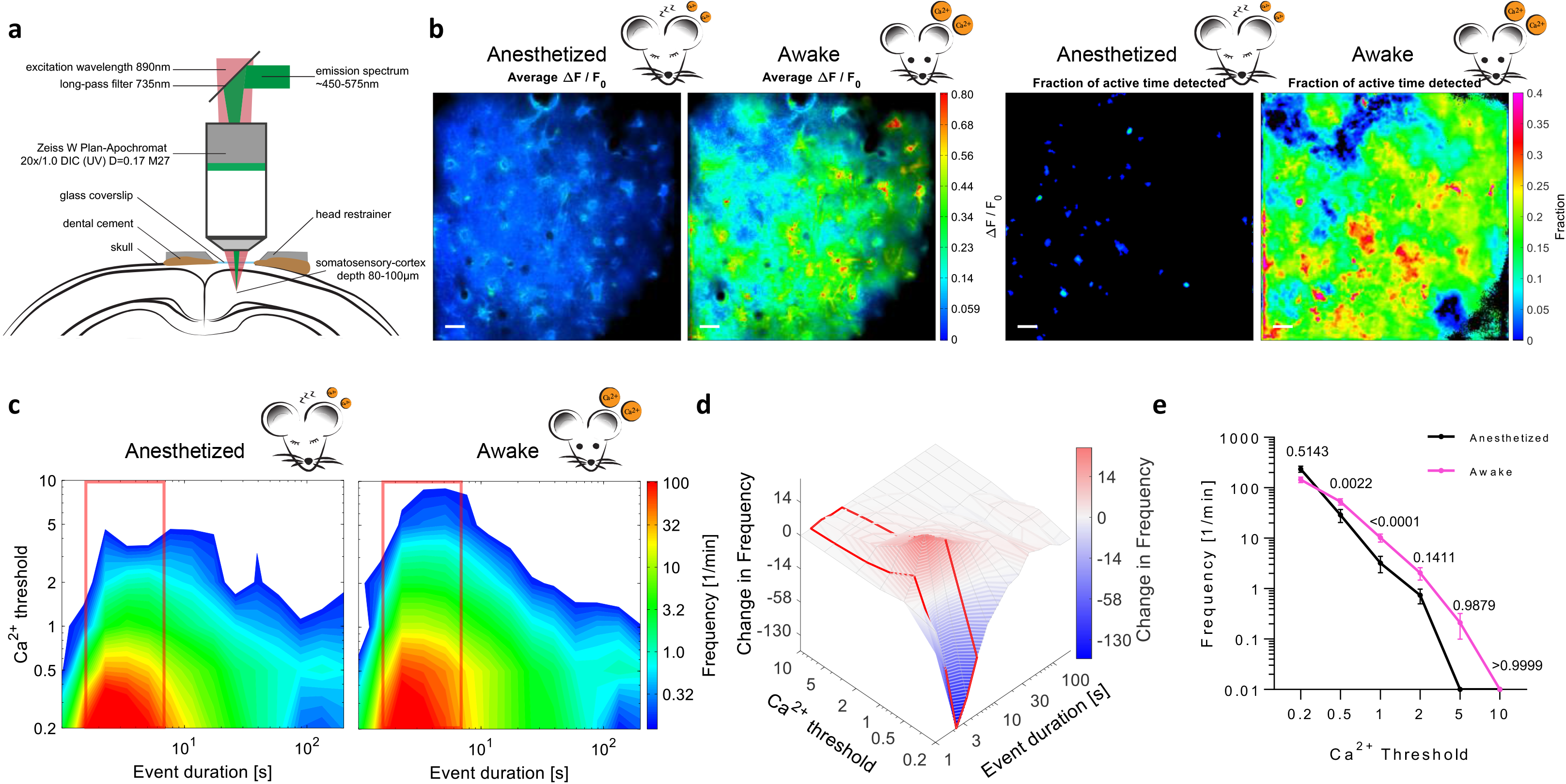
Ca^2+^ event detection strategy MTED is applicable to datasets acquired through *in vivo* cortical imaging. **a**, Experimental scheme of *in vivo* Ca^2+^ imaging through a cortical cranial window. **b**, Astrocytic Ca^2+^ activity in the frontal cortex of a transgenic mouse expressing GCaMP3, represented as the average Ca^2+^ activity ΔF/F_0_ (left) and the fraction of active time (right) in the anesthetized and fully awake mouse. Scale bars = 20 µm. **c**, Data for the Ca^2+^ event duration in anesthetized and awake mice summarized in 2D contour plots. **d**, Frequency differences in Ca^2+^ event duration in anesthetized and awake mice. A gamma of 0.5 was applied to emphasize small changes in frequency. **e**, Statistical evaluation of frequency differences in Ca^2+^ event duration from 1.5 to 7 s. Data show mean and SEM. Two-way ANOVA on arcsine transformed data with Sidak’s multiple comparisons post-hoc test. n = 10 recordings from 3 mice in **c-e**.

Overall, the Ca^2+^ activity in cortical astrocytes observed by *in vivo* imaging showed similar features in the Ca^2+^ pattern as obtained from cultured astrocytes measured at 37°C. The datasets further identify and define the impact of neuronal activity on the Ca^2+^ characteristics of astrocytes (Figure 6c and Figure 5d). Importantly, the only necessary parameters for MTED of Ca^2+^ signals were the thresholds of F_0_ and Ca^2+^. More importantly, our approach robustly detected and comprehensively characterized Ca^2+^ event patterns not only *in vitro* and *in situ* but also *in vivo*.

## Discussion

### Calculating ΔF/F_0_ enables extensive Ca^2+^ activity characterization

In general, with the development of fluorescent indicators of biological activities and metabolites, it has become possible to visualize structural activity and function on a subcellular level. In particular, using GCaMP indicators to monitor the Ca^2+^activity in astrocytes revealed the complexity of these processes, which cannot be described as on or off activity (i.e., below or above a particular threshold). Ca^2+^ event characterization in terms of magnitude, duration, propagation, regional connectivity, and correct event origin provides valuable information about this signaling pathway and is fundamental for understanding the functional role of Ca^2+^activity in astrocytes. In the present study, we developed the MTED algorithm, a novel event detection strategy for time-lapse data obtained by fluorescence microscopy. It is pixel-based and uses multiple thresholds. We chose the latter because signal-to-noise-based strategies for detection of events are prone to underestimating activity in regions with low fluorescence intensity (e.g., low indicator concentrations). This can lead to missed events because the signal amplitudes stay below a single threshold. The multi-threshold approach developed here overcomes this problem. We applied our analysis to data of astrocytic Ca^2+^ signaling as a test case. It enabled us to quantitatively analyze multiple aspects of Ca^2+^ event patterns that evolve in time and space. Indeed, the approach maintained its robustness when weak signals from small astrocytic processes in more intact preparations were analyzed. Characterization of such weak signals is important because those processes are in close proximity with the extracellular matrix, surrounding cells, and synapses, and also represent the origin of the majority of Ca^2+^ events in astrocytes ^17,21^. This robustness is also important for other fluorescent indicators that do not display the large fluorescence changes of modern Ca^2+^ indicators such as many FRET-based sensors.

For all indicators, the relationship between indicator fluorescence and ligand concentration requires careful assessment. In many cases, this relationship is non-linear, and it needs careful consideration to what extent this affects the interpretation of the results. In our study, we used a common measure of fluorescence intensity and its changes (ΔF/F_0_). While this quantification has its drawbacks, see for instance Semyanov et al. 2020^8^, it is straight-forward and abundantly used throughout biomedical research and therefore suitable for testing the analytical approach and its versatility. Since the MTED algorithm is oblivious to the type of data it is working on and does not make specific assumption about it, it can be fed with more quantitative data (e.g. [Ca^2+^] after careful calibration of the imaging setup). Furthermore, it is not limited to Ca^2+^ signaling and could be applied to various fluorescent indicator families and readouts.

### The Ca^2+^ activity pattern in astrocytes is temperature-dependent

Comparing the Ca^2+^ event detection strategies, some predominately describe only fast and regionally restricted event spots, from which they conclude that discussing frequency patterns in restricted active regions is sufficient for the characterization of astrocytic Ca^2+^ activity^16,22^. Other approaches identify a substantial fraction of long-lasting events with regional growth and overlapping patterns, which require the analysis of dynamic regions^17,18^. Here, we observed both types; when examining Ca^2+^ activity at physiological temperature (i.e., 37°C), we observed the first, whereas imaging the same region at RT led to observation of the latter scenario. Our MTED algorithm sufficiently describes both states of this reversible behavior. We revealed that changes in Ca^2+^ event patterns rely on changes in Ca^2+^clearance from the cytosol, which is temperature-dependent^23–25^. A relatively straight-forward explanation is that when the Ca^2+^ uptake is slowed down, Ca^2+^ diffuses to larger volumes, accumulates, and initiates Ca^2+^-induced Ca^2+^ release leading to high magnitude Ca^2+^transients. Blocking the Ca^2+^ uptake by CPA at 37°C indeed switched the Ca^2+^ activity profile to the patterns obtained at 25°C. Remarkably, minor deviations from the species-dependent physiological temperature, such as cooling to 34°C (a temperature typically used in patch-clamp experiments), cause prominent aberrations in the characteristics of Ca^2+^ activity.

Our observation of the temperature-dependent Ca^2+^ activity has broad-ranging consequences for interpreting published findings, as well as for future experimental designs. Deviations between astrocytic Ca^2+^ behavior obtained *in vitro, in situ*, and *in vivo* may be partly explained by different environmental conditions: 25°C for primary cultures, 34°C for organotypic slices, and 37°C for *in vivo* measurements. More importantly, Ca^2+^ activity patterns obtained in our *ex vivo* experiments at 37°C were like the results obtained *in vivo*. Consequently, with advancements in experimental conditions, *in situ* and *in vitro* studies could provide data sets comparable to the *in vivo* situation. This could be beneficial in terms of the 3R (reduce, refine, replace)^26^ and animal welfare. Importantly, the dissection of the temperature dependence of astrocytic Ca^2+^ signaling demonstrates that our analytical approach can be used to compare experimental results across preparations and experimental conditions.

### Cellular environment and state of consciousness shape Ca^2+^ signals in astrocytes

As a second test scenario, we explored how astrocytic Ca^2+^ event patterns are shaped by neuronal activity. We found, for instance, that the occurrence of long-lasting events can be reduced by blocking action potentials in organotypic slices and mixed cultures of astrocytes and neurons with the application of TTX. This adds to previous studies on how neuronal activity modifies astrocytic Ca^2+^ signaling^27^, by identifying the subset of Ca^2+^ events that is impacted. The obtained results suggest that the connectivity between co-cultured neurons and astrocytes is not as dense as in organotypic slice cultures (compare Figure 5d, e and Supplementary Figure 6). An advantage of the presented analysis is the simpler visualization and identification of groups of events that depend on a specific experimental condition, in our example the presence of neurons/neuronal activity. Furthermore, the location and time of this special group of long-lasting Ca^2+^ events can be identified and could potentially be correlated with nearby neuronal structures. More generally, this approach may help with associating spatiotemporal patterns and properties of cellular events reported by fluorescence with biologically relevant mechanisms and conditions.

As a first step towards the latter, we analyzed Ca^2+^ transients of anesthetized and awake animals. Doing so confirmed that Ca^2+^ activity in astrocytes is reduced under anesthetized conditions^28^. Interestingly, this is partly because of a reduction of pronounced Ca^2+^ events with large magnitude whereas the frequency of small magnitude events did not significantly change: an important distinction and refinement. This is a third example of how the MTED analysis can extract important additional information from time-lapse fluorescence microscopy, a type of data set ubiquitous in neurobiology and beyond.

## Methods

### Animals for *in vitro* Ca^2+^ imaging

For all *in vitro* experiments, wildtype animals of both genders from strain C57BL/6J were used. Animals were housed and cared for in accordance to directive 2010/63/EU. Mice were kept in a 14 h light and 10 h dark cycle with lights on starting at 7 am. Animals had *ad libitum* access to food and water and were kept under standard conditions at 22 ± 2 °C RT with 55 ± 5% humidity. Mice were killed by decapitation, and all experiments were conducted according to the recommendations of the European commission.

### Animals for *in vivo* Ca^2+^ imaging

Mice were maintained in the animal facility of the Center for Integrative Physiology and Molecular Medicine (CIPMM, University of Saarland). Astrocyte-specific knock-in GLAST-CreERT2 mice (Slc1a3tm1(cre/ERT2)Mgoe, MGI:3830051)^29^ were crossbred to Rosa26 reporter mice with GCaMP3 expression (Gt(ROSA)26Sortm1(CAG-GCaMP3)Dbe, MGI: 5659933)^30^. Imaging sessions were performed at 8-10 weeks of age. Mouse administration was managed via the PyRAT database (Python based Relational Animal Tracking) from Scionics Computer Innovation GmbH (Dresden, Germany). Animals were kept and bred in strict accordance with the recommendations to European and German guidelines for the welfare of experimental animals. Animal experiments were approved by the Saarland state’s “Landesamt für Gesundheit und Verbraucherschutz” in Saarbrücken/Germany (license numbers: 71/2013 and 36/2016).

### Primary hippocampal astrocyte cultures

Primary astrocyte cell cultures were prepared according to a previously described protocol^31^ with slight modifications: Hippocampi were isolated from brains of neonatal mice between P1-3 and cells were seeded after dissociation at a density of 5×10^4^ cells per 12 mm glass coverslip for microscopy in 500 µl plating medium (49 ml MEM, 1 ml B-

27 supplement, 500 µl sodium pyruvate, 500 µl L-Glutamine, 50 µl Penicillin-Streptomycin; all Thermo Fisher Scientific Inc., Waltham, USA). On DIV3 the entire plating medium was replaced with 1 ml maintenance medium (49 ml Neurobasal-A, 1 ml B-27 supplement, 500 µl L-Glutamine, 50 µl Penicillin-Streptomycin; all Thermo Fisher Scientific Inc., Waltham, USA). On DIV11, ½ of the medium was exchanged with prewarmed maintenance medium prior to infection of the cells with of 0.1 µl AAV-mGFAP-GCaMP6s (3.7 x 10^9^vg/µl) and AAV-mGFAP-tdTomato (1 x 10^7^ vg/µl). Astrocytes were maintained at 37 °C in a humidified incubator in a 5% CO_2_atmosphere used for experiments between DIV14-17. Cells were transferred to a prewarmed recording chamber for microscopy and kept in a balanced salt solution (BSS), which was adjusted to pH 7.4 and 290 mOsm with glucose, containing 115 mM NaCl, 5.4 mM KCl, 1 mM MgCl_2_, 2 mM CaCl_2_ and 20 mM HEPES.

### Organotypic slice cultures

Organotypic slice cultures were prepared after an adapted protocol from Kobe *et al*.^32^. Briefly, mice were decapitated at P6 under sterile conditions and the isolated hippocampus was placed in ice-cold oxygenized slice medium in a 60 mm dish for 30 min. 350 µm thick slices were prepared with McIlwain Tissue Chopper (Mickle, Surrey, UK) and separated with a needle to select 2-4 slices with complete hippocampal structures. Selected slices were transferred onto Millicell filter inserts (#PICM03050, Merck, Darmstadt, Germany) in a 6-well plate containing 1 ml slice maintenance medium (50% MEM, 25% Hanks’ balanced salt solution, 25% horse serum, and 2 mM glutamine at pH 7.3). Excess liquid around the slice was removed and cells were subsequently infected by application of 0.2 µl AAV-mGFAP-GCaMP6s (3.7 x 10^9^ vg/µl) into the medium. Slices were kept in a humidified atmosphere (5% CO_2_, 37°C) with ½ of the medium being exchanged on DIV2, DIV4 and DIV6. Ca^2+^ imaging was conducted at DIV5-7.

### Cranial window surgery for *in vivo* two-photon imaging

During surgical procedures, animals were kept on heating pads and eyes were covered with Bepanthen ointment (Bayer, Leverkusen, Germany). Anesthesia was induced with a mixture of 5 % isoflurane, 47.5 % O_2_ (0.6 l/ min) and 47.5 % N_2_O (0.4 l/ min) and maintained with 2 % isoflurane (Harvard Apparatus anesthetic vaporizer, Harvard, Holliston, USA). A standard craniotomy^33^of 3 mm in diameter was performed over the somatosensory cortex (2 mm posterior and 1.5 mm lateral to bregma). The craniotomy was sealed with a glass coverslip (Glaswarenfabrik Karl Hecht, Sondheim, Germany; #1.5 thickness code) and fixed with dental cement (RelyX®, 3M ESPE, Seefeld, Germany). Subsequently, a metal holder for head restraining (5 mm diameter) was applied and fixed to the skull with dental cement. After surgery, the animals were kept on the heating pad until complete recovery. Post-operative treatment consisted of buprenorphine (3 mg/kg, s.c.) and dexamethasone (0.2 mg/kg, i.p.), for three consecutive days. Recovery was assessed by body weight and mouse grimace scale. After five to seven days the first imaging session was performed.

In preparation for Ca^2+^ imaging, animals were habituated before the first imaging session according to adapted protocols without water restriction from Guo *et al*.^34^ and Kislin et al.^35^ The animals were head-fixed with a custom-designed head-restrainer, 3D-printed using stainless steel. During imaging, anesthesia was applied using a custom-made, magnetically attachable anesthesia mask. Each field of view (FOV) was imaged twice: first in anesthetized, then in awake state. During imaging in anesthetized state, isoflurane concentration was kept at 1.5 %, and flow of O_2_ and N_2_O was set to 0.6 l/min and 0.4l /min, respectively. Before awake state imaging, isoflurane and other gases were switched off and it was verified that the animals were fully awake. The selected FOVs for Ca^2+^ imaging were located in the somatosensory cortex, 80 – 100 μm beneath the dura. Each FOV was recorded for 5 min to investigate the Ca^2+^ signals. The total duration of one imaging session ranged between 30-60 min per animal. After imaging, animals were kept on a heating pad at 37°C until they recovered completely, additionally Fresubin (Fresenius Kabi, Bad Homburg, Germany) was provided *ad libitum*.

### Reagents

Tetrodotoxin citrate (TTX; #Asc-055; Ascent Scientific, Princeton, NJ) was used at a concentration of 10 nM to block neuronal activity and was applied several minutes prior to imaging. The Ca^2+^-ATPase inhibitor Cyclopiazonic acid (CPA; #120300, Abcam, Cambridge, UK) was applied at a concentration of 10 µM at least 10 min before the measurements.

### Microscopy

Ca^2+^ imaging *in vitro* and *in situ* was conducted on an upright Andor Spinning Disc microscope (Oxford Instruments, Belfast, Northern Ireland) equipped with a CSU-X1 (Yokogawa, Musashino, Japan) using filter cubes (537/26 nm) for full frame imaging of GCaMP6s or split filter cubes (609/54) for simultaneous imaging of GCaMP6s and tdTomato. Cells were recorded for 10 min with 5 frames/s using excitation wavelength 488 nm (GCaMP6s) and 561 nm (tdTomato). The temperature of the BSS for measurement was controlled by a custom-built heating device and additionally supervised with an external thermometer. To achieve thermal stability and avoid artefacts during recordings the objective, stage and chamber were all heated to the desired temperature. *In vivo* Ca^2+^ imaging was conducted in anaesthetized (1.5% isoflurane) and awake head-fixed mice through a cortical cranial window in the prefrontal cortex using two-photon excitation laser scanning microscopy (TPE-LSM). The custom-built microscope was equipped with a resonant scanner (RESSCAN-MOM, Sutter instrument, Novato, USA) and a 20x water-immersion objective (W Plan-Apochromat 20x/1.0 DIC D=0.17; Zeiss, Jena, Germany). Images were acquired with a frame rate of 30 Hz and a 10 Hz frame-averaging factor, resulting in an effective acquisition rate of 3 Hz. To minimize photo-damage, the excitation laser power was kept at a minimum for a sufficient signal-to-noise ratio (<40 mW at 60 ns pixel dwell time). Laser wavelength was set to 890 nm (Chameleon Ultra II, Ti:Sapphire Laser; Coherent, Santa Clara, USA). The emitted light was detected by a photomultiplier tube (R6357; Hamamatsu, Hamamatsu, Japan) and pre-amplified (DHPCA-100, Femto, Berlin, Germany). Digitizer (NI-5734) and control hardware (NI-6341) was housed in a PXIe (1082) chassis, connected to a control-PC via a high bandwidth PXIe-PCIe8398 interface (NI, Austin, USA). Scanning and image acquisition were controlled by ScanImage (SI 5.6R1)^36^.

### Data processing

Data were processed using Matlab. *In vitro* data obtained by confocal spinning disk microscopy were denoised using VBM3D^37^ and low intensity TPE-LSM data by SURE-LET. An automated data offset control was obtained by analyzing the data intensity histogram. The *F*_0_ algorithm is based on ‘moving window’ filter functions, where the mean and variance of *F* are used as weighting functions for *F* to identify low intensities. The ‘moving window’ sizes for filtering are input parameters, dependent on acquisition settings and the temporal profile of expected signal changes. The MTED algorithm uses *F* and *F*_0_ or *F*_*R*_ and applied various Ca^2+^ thresholds to Δ*F*/*F*_0_ or Δ*F*/*F*_*R*_, respectively. To reject false-positive pixels, we applied an appropriate *F*_0_ threshold to eliminate background signals, such as readout noise. Next, basic morphological gray-scale operations (opening and closing) were performed to close holes and remove small isolated peaks, followed by Gaussian smoothing of the Ca^2+^ signals. This was used as a weighting function. Groups of Ca^2+^-positive regions were then identified based on their spatio-temporal connectivity. Small groups not exhibiting a minimum size and duration were rejected (see Supplementary Movie 3). The time dependence of a detected event was stored for visualization and further analysis. The Ca^2+^ event detection was repeated for various Ca^2+^ thresholds. For practical reasons, we use logarithmic-like spaced threshold levels, such as [0.2, 0.5, 1, 2, 5, 10] fold change in *F*.

## Data availability

The data sets generated and analyzed in this study are available from the corresponding author upon reasonable request (approximately 3.5 TB).

## Code availability

The Matlab code developed in this study is available from the corresponding author upon reasonable request.

## Acknowledgement

This study was supported by the German Research Foundation (DFG) (PO732 to EGP, ZE994/2 to AZ, SFB1089 B03, SPP1757 HE6949/1 and HE6949/3 to CH, and by SFB 894 A12 and SPP 1757 KI 503/12-2 to FK). This manuscript is part of the PhD thesis of FEM.

## Author Contributions

FEM, GS, VC, LCC and LS conducted laboratory work; FK, CH, EGP and AZ shaped the experimental outline and supervised the project; GS, VC and AZ wrote the algorithm; FEM and AZ wrote the initial manuscript which was then contributed to by all co-authors.

## Competing Interests statement

The authors declare no conflicts of interest.

## Supplementary Figures and Movies

**Supplementary Figure 1:**
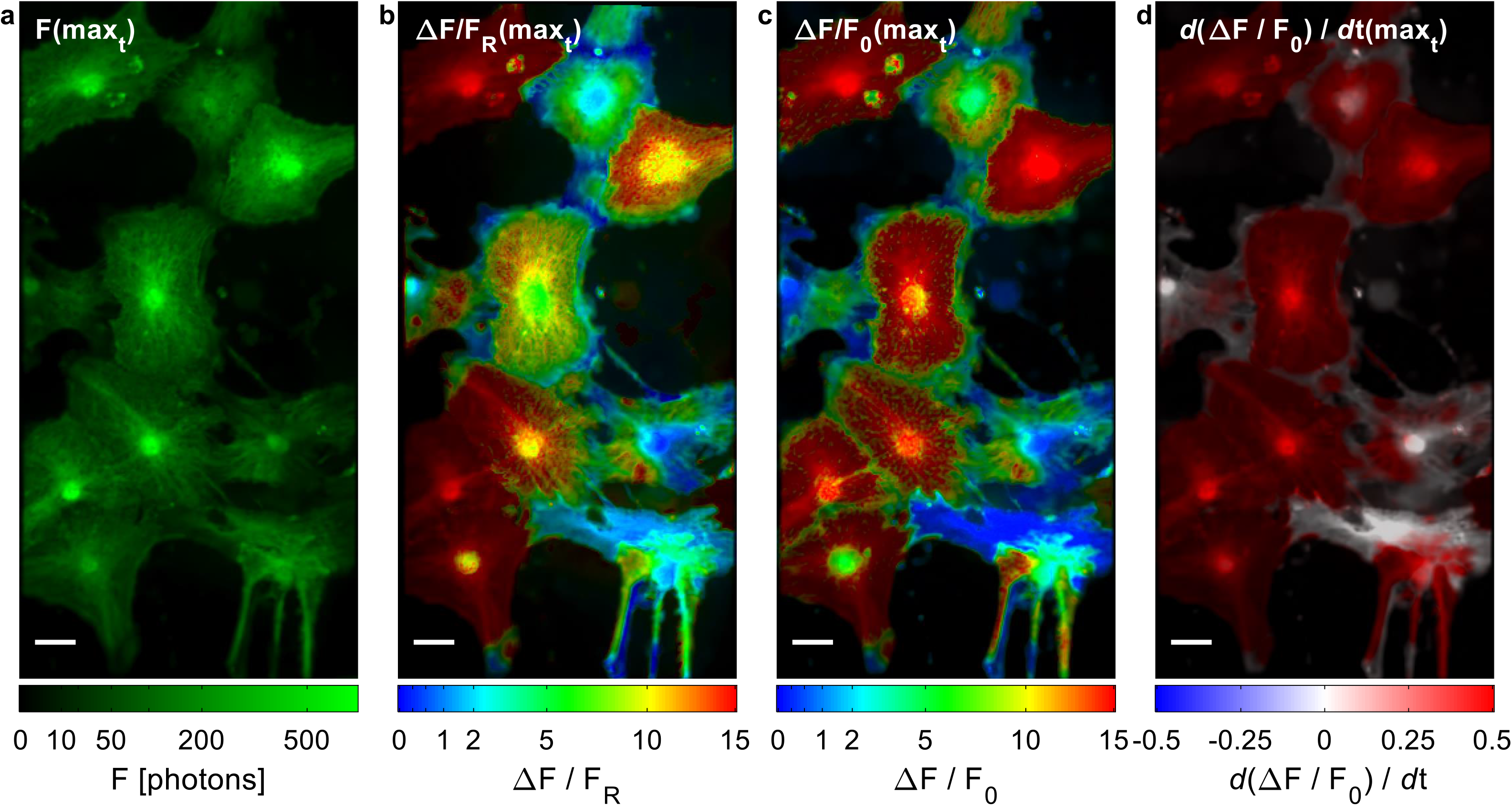
Ca^2+^ activity of cultured astrocytes comparing F, ΔF/F_R_ and ΔF/F_0_. **a**, Maximum projection of F over time to identify astrocyte morphology. The image provides no information about Ca^2+^ activity. **b-c**, Maximum projection of ΔF/F_R_ and ΔF/F_0_ over time, scale bar = 10 µm, color code according to Figure 1c-d. **b**, The color code reveals regions of high (red) and of low (blue) maximal Ca^2+^ activity, while the maximal change in Ca^2+^ seems to vary from cell to cell. However, this is most likely caused by different expression levels of the Ca^2+^ indicator GCaMP6s and tdTomato, which acts as ruler. Cell specific differences in the basal [Ca^2+^] cannot be excluded. To this end, a cell specific scaling factor would be required to overcome misinterpretation. **c**, The concept of ΔF/F_0_ does not require a cell-specific scaling factor. However, it relies on the accuracy of F_0_, which is obtained in a pixel-based manner. Differences in basal [Ca^2+^] cannot be identified. **d**, The time-maximum projection of temporal changes in Ca^2+^ (compare Figure 1d) is a direct outcome of the ΔF/F_0_ data. The red color indicates a strong increase of Ca^2+^, e.g. all over the central cell.

**Supplementary Figure 2:**
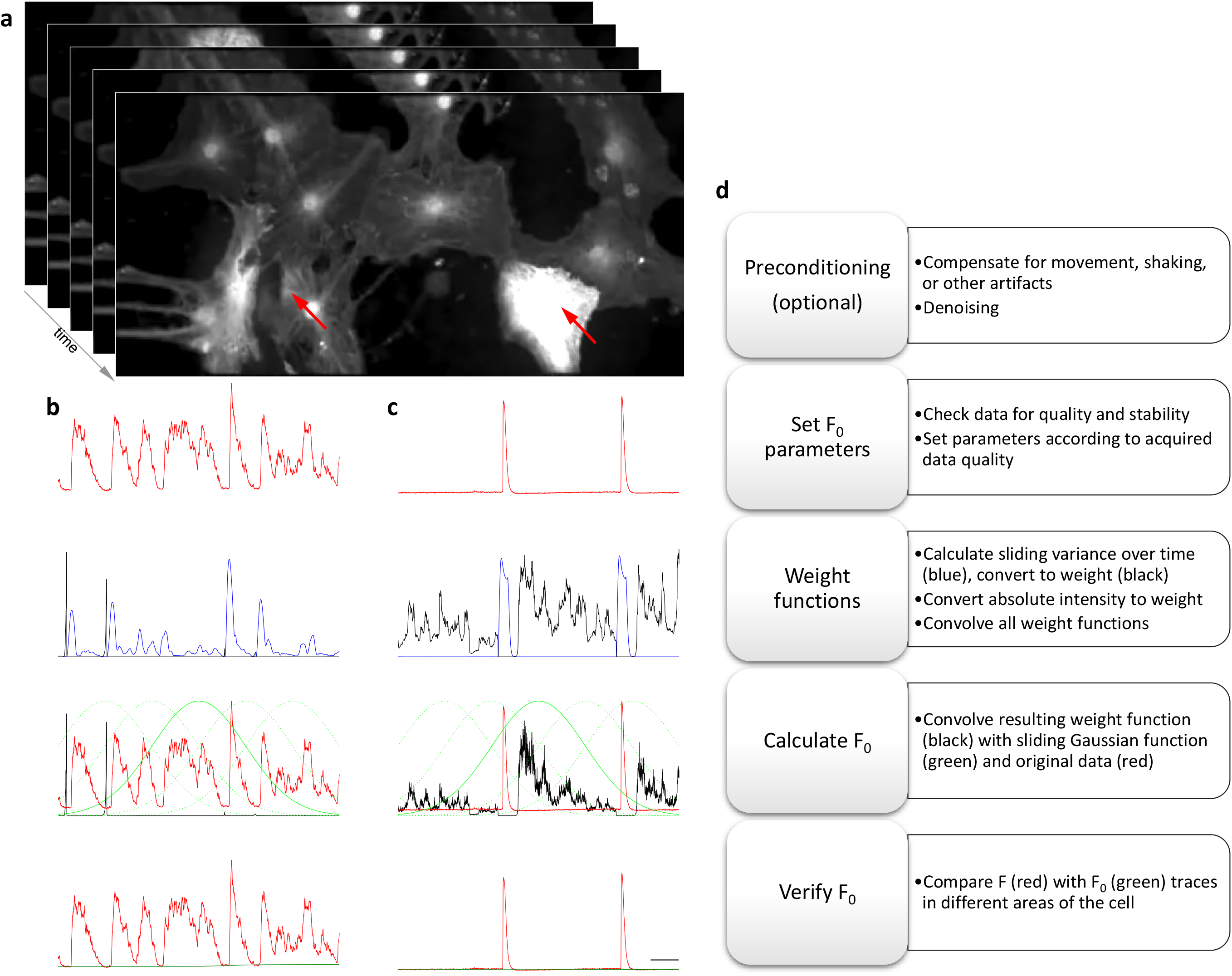
Automatic F_0_ estimation from intrinsic fluctuation analysis of the fluorescence signal of Ca^2+^ indicators. **a**, The preprocessed image sequence of the Ca^2+^ indicator GCaMP6s fluorescence signal is used to estimate F_0_ for each pixel independently, no neighboring information is used. **b-c**, The time traces of two individual pixels, highlighted by the two red arrows in a. **d**, Description of the specific calculation procedure and the color code used in b-c.

**Supplementary Figure 3:**
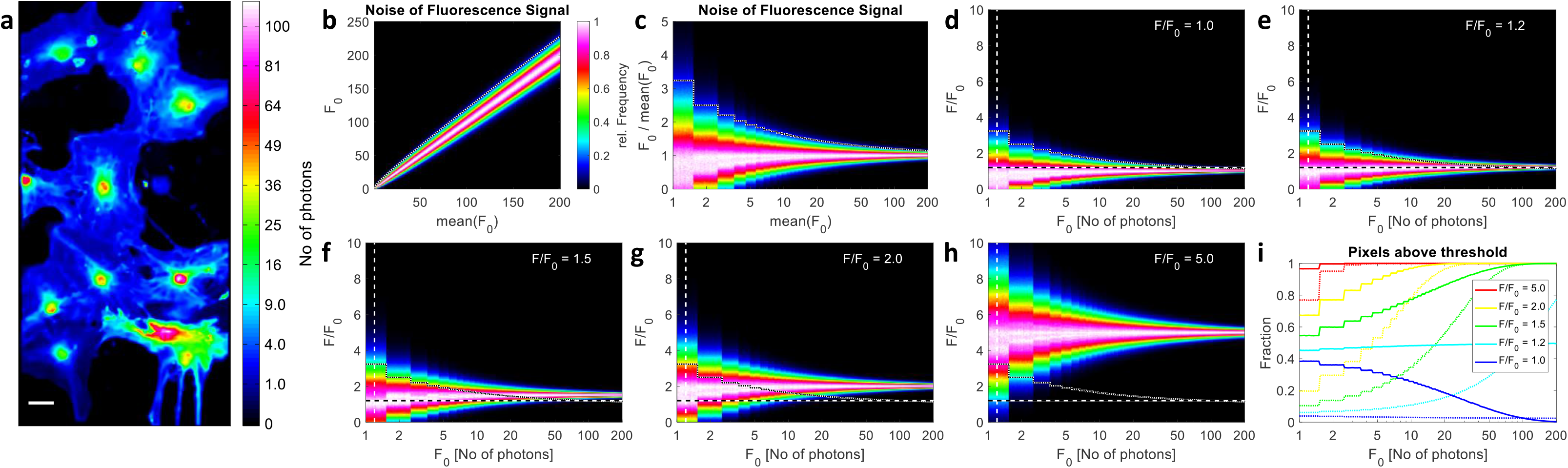
Simulations of thresholding accuracy according to signal strength. **a**, The signal strength of F_0_ from Figure 1, shown as number of photons detected per pixel and each timepoint, illustrates the expected accuracy of F/F_0_. **b**, From noise simulations (images with 1000 x 1000 pixel, intensities from 1 to 200 photons, applying Poisson noise overlaid by a gaussian detector noise, sigma 0.5) the expected noise was estimated, the mean +2x standard deviation is shown as a dashed white line and depicts a common concept to set a threshold and distinguish signal from background. **c**, Display of the data from b as F/F_0_ reveals the conceptional problem of using the general threshold 2x standard deviation above noise level. As for small F_0_ values only very strong Ca^2+^ changes lie above threshold, small Ca^2+^ changes are consequently not detected when F_0_ is small, which is typically the case in small processes of astrocytes. **d-h**, The signal distribution as a function of F_0_ for signals F/F_0_ = 1, 1.2, 1.5, 2, 5, which represent ΔF/F_0_ = 0, 0.2, 0.5, 1, 4. The horizontal dashed line represents the 0.2 threshold for F/F_0_, i.e. for Ca^2+^ change, the vertical for F_0_. The dotted black & white lines (compare b-c) show the threshold with 2x standard deviation above F_0_ noise level. **i**, From the simulations the fraction of pixels above threshold could be obtained for both concepts, the fixed threshold (solid lines) and the threshold with 2x standard deviation above F_0_ (dotted lines). As expected, the fixed threshold concept provides a much higher chance to identify small Ca^2+^ changes at low F_0_. The relatively high number of false positive detected pixel (blue solid line) requires additional processing steps (compare Figure 2).

**Supplementary Figure 4:**
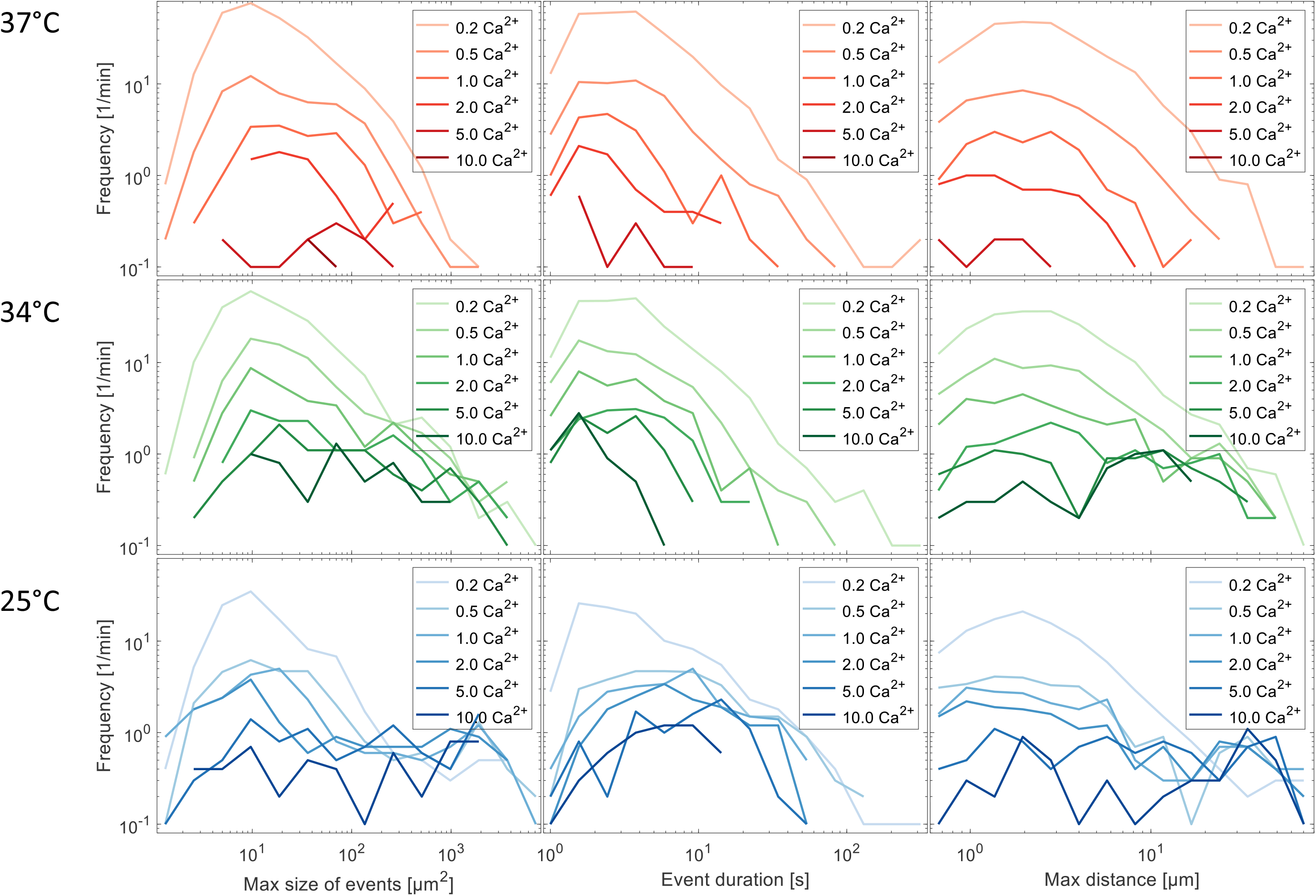
Statistical characterization of Ca^2+^ activity in astrocytes at different temperatures. To better illustrate the origin of the 2D histogram plots in Figure 3c, the data are shown here as line histograms, where the color indicates the environmental temperature (blue: 25°C, green: 34°C, red: 37°C) and the color opacity reflects the Ca^2+^ threshold, as indicated in the figure legends.

**Supplementary Figure 5:**
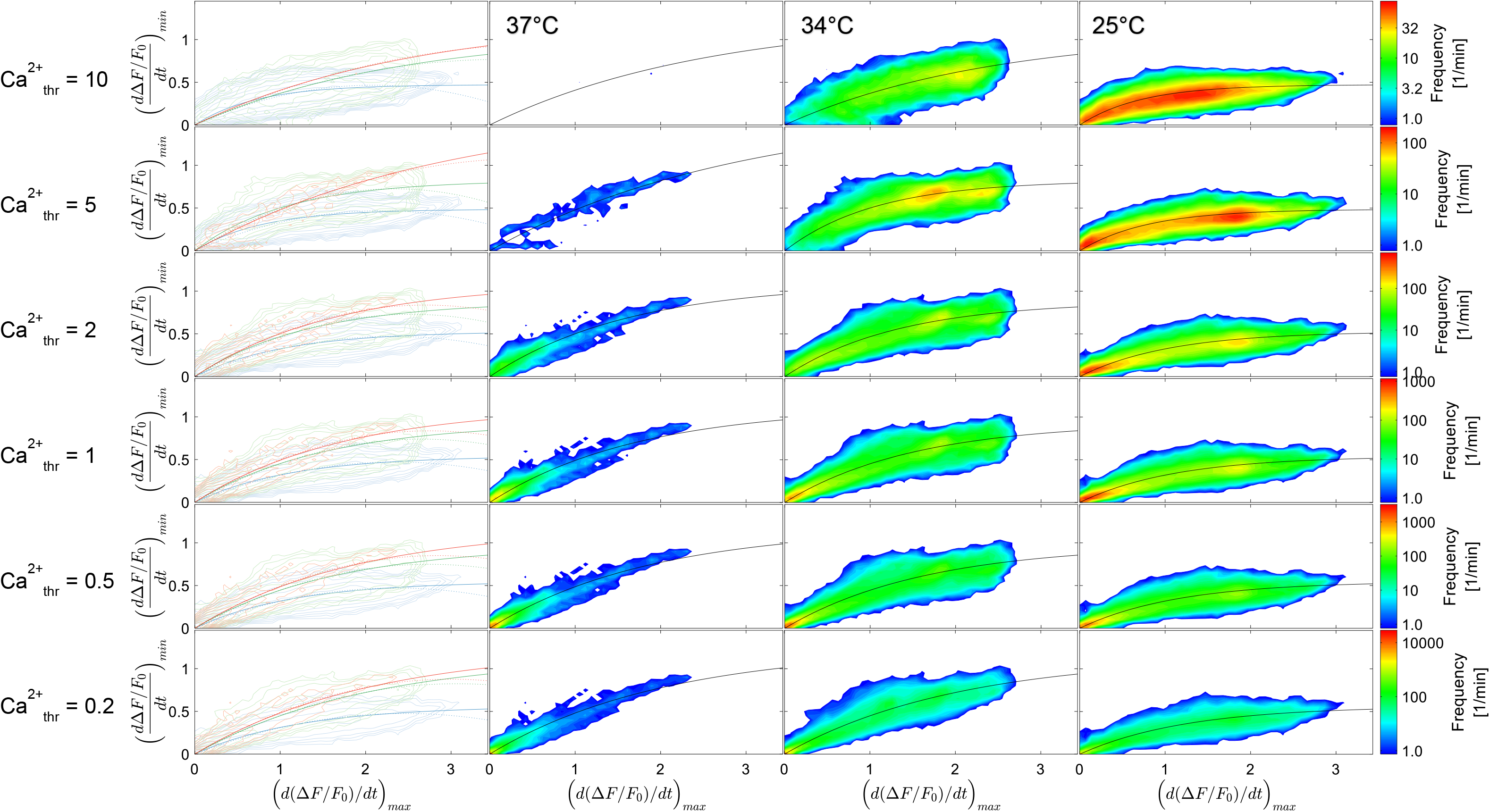
Relation between maximal Ca^2+^ increase and maximal Ca^2+^ decrease for different thresholds and temperatures depicted as 2D histograms. The maximum and the minimum of the first derivative of ΔF/F_0_ was plotted for each pixel as 2D histogram. To illustrate the relation between both parameters, the data were fitted by the exponential function y = a · (1 − e^−bx^) and shown as black line for the individual temperature plots and color coded in the overlaid contour plots. Whereas at low temperatures high maximal positive slopes were found, which could be interpreted as strong Ca^2+^ releases, the same pixel shows only a moderate minimal negative slope, which does not increase linearly with increasing positive change. Since Ca^2+^ release and uptake are competing processes, at higher environmental temperatures the maximum Ca^2+^ increase is reduced, which should not be interpreted as a reduced Ca^2+^ release rate but indicate a competing Ca^2+^ uptake. The left row overlays all three temperature patterns for the specific thresholds.

**Supplementary Figure 6:**
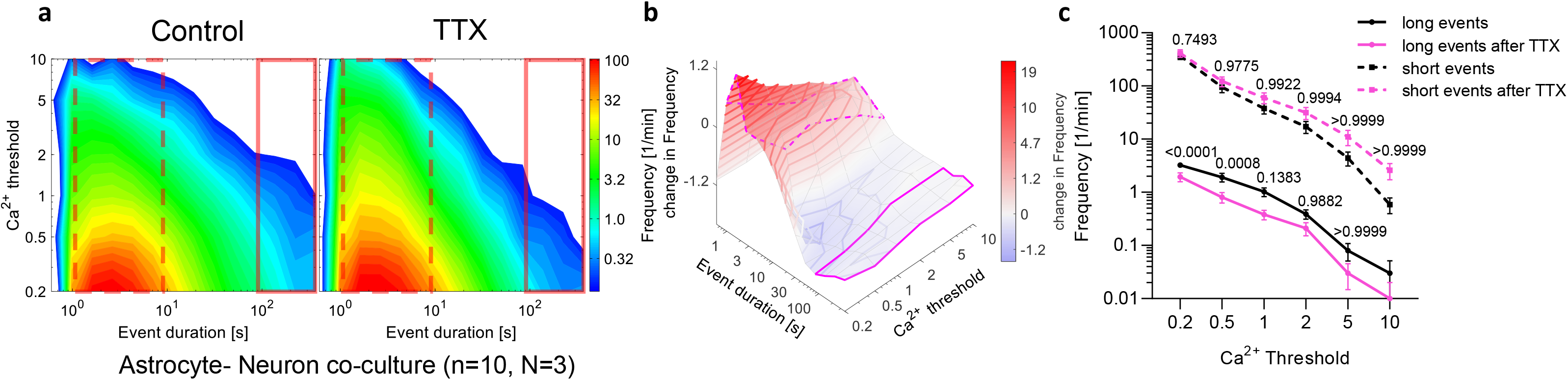
Blocking of neuronal activity in astrocyte-neuron co-cultures. **a**, Summarized 2D contour plots of the event duration pattern without and with application of TTX. **b**, Frequency differences in Ca^2+^ event duration without and in presence of TTX. **c**, Statistical evaluation shows the frequency reduction of long lasting Ca^2+^ events (> 90 s), while short events (1 - 10 s) are not affected.

**Supplementary Figure 7:**
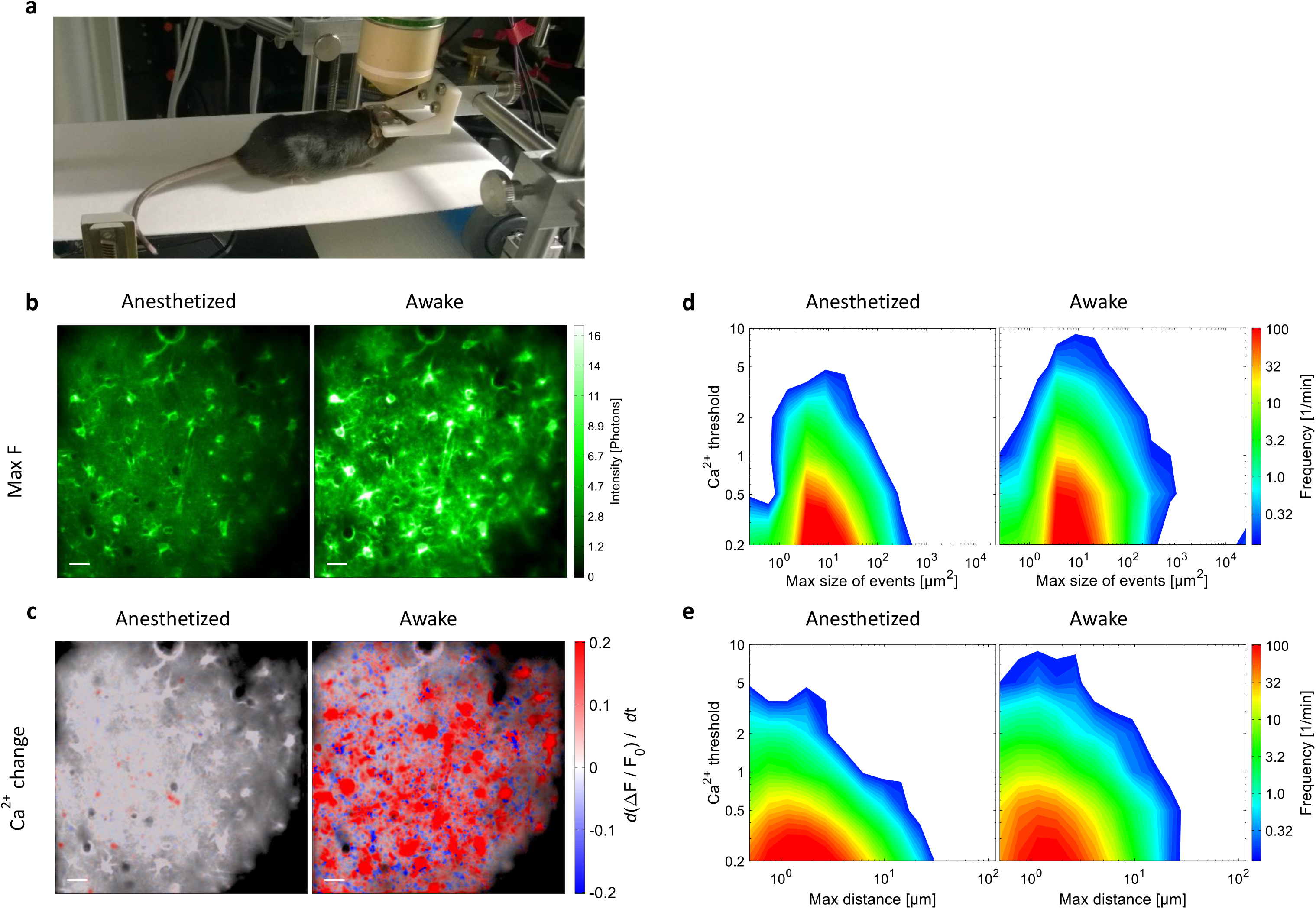
Ca^2+^ activity *in vivo* of anesthetized and awake transgenic mice. **a**, Setup for *in vivo* Ca^2+^ imaging. **b**, Maximum fluorescence signal F shows higher Ca^2+^ activity in the same region when mice are awake. Scale bar = 20 µm. **c**, Visualization of changes in Ca^2+^, which are more abundant, prominent, and extended in the awake state. Scale bar = 20 µm. **d-e**, 2D contour plots show the maximum size of Ca^2+^ events (d) and the maximum distance (e), which are both reduced when animals were anesthetized as compared to when mice were awake during measurements. n = 10 recordings from N = 3 mice.

**Supplementary Table 1:**
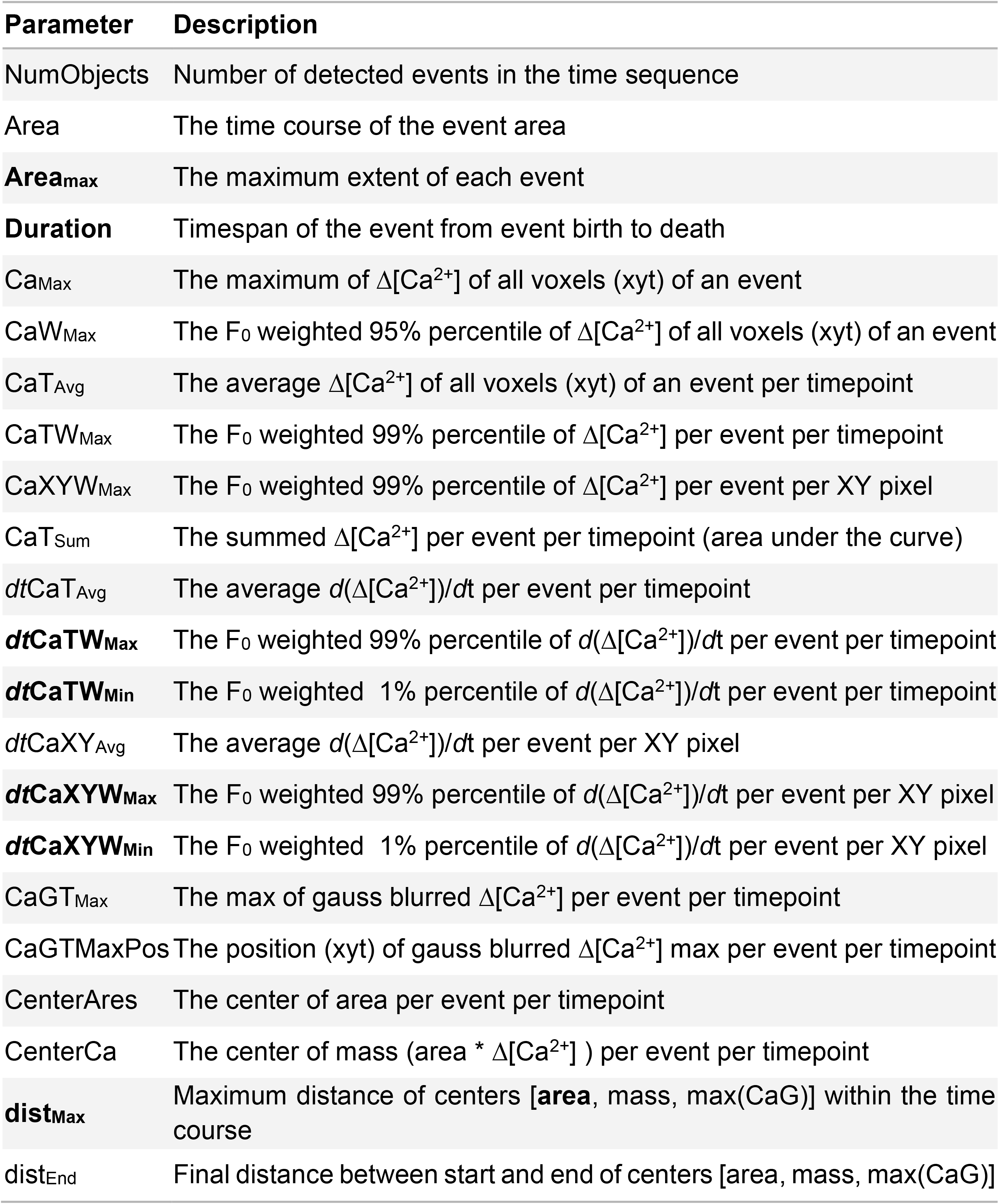
Overview of calculated parameters in a standard evaluation. The parameters are obtained for each Ca^2+^ threshold level. Parameters in bold are used in this study.

| Parameter | Description |
| --- | --- |
| NumObjects | Number of detected events in the time sequence |
| Area | The time course of the event area |
| **Area<sub>max</sub>** | The maximum extent of each event |
| **Duration** | Timespan of the event from event birth to death |
| Ca<sub>Max</sub> | The maximum of $\Delta[\text{Ca}^{2+}]$ of all voxels (xyt) of an event |
| CaW<sub>Max</sub> | The $F_0$ weighted 95% percentile of $\Delta[\text{Ca}^{2+}]$ of all voxels (xyt) of an event |
| CaT<sub>Avg</sub> | The average $\Delta[\text{Ca}^{2+}]$ of all voxels (xyt) of an event per timepoint |
| CaTW<sub>Max</sub> | The $F_0$ weighted 99% percentile of $\Delta[\text{Ca}^{2+}]$ per event per timepoint |
| CaXYW<sub>Max</sub> | The $F_0$ weighted 99% percentile of $\Delta[\text{Ca}^{2+}]$ per event per XY pixel |
| CaT<sub>Sum</sub> | The summed $\Delta[\text{Ca}^{2+}]$ per event per timepoint (area under the curve) |
| dtCaT<sub>Avg</sub> | The average $d(\Delta[\text{Ca}^{2+}])/dt$ per event per timepoint |
| **dtCaTW<sub>Max</sub>** | The $F_0$ weighted 99% percentile of $d(\Delta[\text{Ca}^{2+}])/dt$ per event per timepoint |
| **dtCaTW<sub>Min</sub>** | The $F_0$ weighted 1% percentile of $d(\Delta[\text{Ca}^{2+}])/dt$ per event per timepoint |
| dtCaXY<sub>Avg</sub> | The average $d(\Delta[\text{Ca}^{2+}])/dt$ per event per XY pixel |
| **dtCaXYW<sub>Max</sub>** | The $F_0$ weighted 99% percentile of $d(\Delta[\text{Ca}^{2+}])/dt$ per event per XY pixel |
| **dtCaXYW<sub>Min</sub>** | The $F_0$ weighted 1% percentile of $d(\Delta[\text{Ca}^{2+}])/dt$ per event per XY pixel |
| CaGT<sub>Max</sub> | The max of gauss blurred $\Delta[\text{Ca}^{2+}]$ per event per timepoint |
| CaGTMaxPos | The position (xyt) of gauss blurred $\Delta[\text{Ca}^{2+}]$ max per event per timepoint |
| CenterAres | The center of area per event per timepoint |
| CenterCa | The center of mass (area * $\Delta[\text{Ca}^{2+}]$) per event per timepoint |
| **dist<sub>Max</sub>** | Maximum distance of centers [**area**, mass, max(CaG)] within the time course |
| dist<sub>End</sub> | Final distance between start and end of centers [area, mass, max(CaG)] |

## Supplementary Movie Captions

**Supplementary Movie 1:** Ca^2+^ activity of cultured astrocytes comparing F, ΔF/F_R_ and ΔF/F_0_ (related to Suppl. Figure 1).

**a**, Time sequence of the denoised Ca^2+^ indicator signal F. The temporal change of F provides already an idea of the underlying Ca^2+^ activity. **b-c**, Time sequence of ΔF/F_R_ and ΔF/F_0_. Scale bar = 10 µm, color code according to Figure 1c-d. **d** The color coded temporal change in Ca^2+^ obtained from ΔF/F_0_(*d*(ΔF/F_0_)/*d*t(max_t_)). Red colors depict an increase in Ca^2+^ levels, blue colors a decrease, respectively. Scale bar = 10 µm, color code according to Figure 1d.

**Supplementary Movie 2:** Visualization of the functionality of F_0_ calculation.

The time sequence from Figure 1 and Suppl. Figure 1 was used to visualize the time course of F and F_0_. **Left**, Color coded sequence of ΔF/F_0_, including a traveling cross depicting the position of the time profile of F and F_0_ shown in the right graph. **Right**, Time profile of F and F_0_ of the xy position depicted by the cross in the left movie sequence.

**Supplementary Movie 3:** Individual steps of the Ca^2+^ event detection algorithm.

The time sequence from Figure 1b-f, was used to visualize the individual steps of the Ca^2+^ event detection algorithm (see Figure 2a). **a**, Binary image sequences after applying Ca^2+^ threshold 0.2. **b**, Image sequences after applying F_0_ threshold. Red areas are removed pixels due to the threshold. White areas are remaining regions. **c**, Red areas are removed pixels due to morphological filtering, such as applying stability criteria and erosion. White areas are remaining regions. **d**, Remaining regions before removing small groups, neighboring pixels have the same color. **e**, Final outcome from the Ca^2+^ event detection algorithm after removing small groups.

**Supplementary Movie 4:** Outcome of the Ca^2+^ event detection algorithm.

The time sequence from Figure 1 was used to visualize the outcome of the Ca^2+^event detection algorithm. **a**, Color coded ΔF/F_0_ presented according to Suppl. Movie 1c. **b**, Color-coded event regions for all Ca^2+^thresholds applied. **c**, Detected events overlaid as contours going from light to dark magenta for low to high Ca^2+^ thresholds, respectively.

**Supplementary Movie 5:** Time sequences of the same field of view at different temperatures.

Primary cultures of hippocampal astrocytes were cooled down from 37°C to 25°C and heated up again to 37°C. The recognized astrocytic Ca^2+^ activity is overlaid as contours (compare Figure 2b,c, Suppl. Movie 4).

**Supplementary Movie 6:** 3D contour plots of detected events at the three different temperatures.

Video sequence of detected events depicted as a one-minute 3D (xyt) contour (see Figure 3b), in which darker colors represent higher ΔCa^2+^.This illustrates the differences in Ca^2+^ activity with environmental temperature.

**Supplementary Movie 7:** Generation of 2D histograms for statistical parameters from frequency plots at different Ca^2+^ thresholds.

In this video sequence a step-by-step illustration is provided how statistical parameters are visualized as 2D frequency histograms (see Figures 3c, 4c, 5c and d, 6c, Suppl. Figures 6a, 7d and e). Here, the event duration is shown exemplarily.

**Supplementary Movie 8:** Astrocytic Ca^2+^ activity in the frontal cortex of a transgenic mouse in anesthetized state expressing GCaMP3.

Video sequence of the denoised GCaMP3 fluorescence signal F. The color coded Ca^2+^ signal ΔF/F_0_ and the same video sequence overlaid by the detected events, depicted as contours going from light to dark magenta for low to high Ca^2+^ thresholds, respectively.

**Supplementary Movie 9:** Astrocytic Ca^2+^ activity in the frontal cortex of a transgenic mouse in awake state expressing GCaMP3.

Video sequence of the denoised GCaMP3 fluorescence signal F. The color coded Ca^2+^ signal ΔF/F_0_ and the same video sequence overlaid by the detected events, depicted as contours going from light to dark magenta for low to high Ca^2+^ thresholds, respectively.

## Notes

### Competing Interest Statement

The authors have declared no competing interest.

## References

1. Chai, H. et al. Neural circuit-specialized astrocytes: transcriptomic, proteomic, morphological and functional evidence. Neuron 95, 531-549.e9 (2017).

2. Panatier, A. et al. Astrocytes Are Endogenous Regulators of Basal Transmission at Central Synapses. Cell 146, 785–798 (2011).

3. Araque, A. et al. Gliotransmitters Travel in Time and Space. Neuron 81, 728–739 (2014).

4. Bonansco, C. et al. Glutamate released spontaneously from astrocytes sets the threshold for synaptic plasticity. European Journal of Neuroscience 33, 1483–1492 (2011).

5. Henneberger, C., Papouin, T., Oliet, S. H. R. & Rusakov, D. A. Long term potentiation depends on release of D-serine from astrocytes. Nature 463, 232–236 (2010).

6. Porter, J. T. & Mccarthy, K. D. Astrocytic neurotransmitter receptors in situ and in vivo. Progress in Neurobiology 51, 439–455 (1997).

7. Volterra, A. & Meldolesi, J. Astrocytes, from brain glue to communication elements: the revolution continues. Nat Rev Neurosci 6, 626–640 (2005).

8. Semyanov, A., Henneberger, C. & Agarwal, A. Making sense of astrocytic calcium signals — from acquisition to interpretation. Nature Reviews Neuroscience 21, 551– 564 (2020).

9. Nett, W. J., Oloff, S. H. & McCarthy, K. D. Hippocampal Astrocytes In Situ Exhibit Calcium Oscillations That Occur Independent of Neuronal Activity. Journal of Neurophysiology 87, 528–537 (2002).

10. Volterra, A., Liaudet, N. & Savtchouk, I. Astrocyte Ca2+ signalling: an unexpected complexity. Nat Rev Neurosci 15, 327–335 (2014).

11. Meldolesi, J. The development of Ca2+ indicators: a breakthrough in pharmacological research. Trends in Pharmacological Sciences 25, 172–174 (2004).

12. Pérez Koldenkova, V. & Nagai, T. Genetically encoded Ca2+ indicators: Properties and evaluation. Biochimica et Biophysica Acta (BBA) - Molecular Cell Research 1833, 1787–1797 (2013).

13. Horikawa, K. Recent progress in the development of genetically encoded Ca2+ indicators. J. Med. Invest. 62, 24–28 (2015).

14. Venugopal, S., Srinivasan, R. & Khakh, B. S. GECIquant: Semi-automated Detection and Quantification of Astrocyte Intracellular Ca2+ Signals Monitored with GCaMP6f. in Computational Glioscience (eds. De Pittà, M. & Berry, H.) 455–470 (Springer International Publishing, 2019). doi:10.1007/978-3-030-00817-8_17.

15. Castro, M. A. D. et al. Local Ca 2+ detection and modulation of synaptic release by astrocytes. Nat Neurosci 14, 1276–1284 (2011).

16. Agarwal, A. et al. Transient Opening of the Mitochondrial Permeability Transition Pore Induces Microdomain Calcium Transients in Astrocyte Processes. Neuron 93, 587-605.e7 (2017).

17. Wu, Y.-W. et al. Morphological profile determines the frequency of spontaneous calcium events in astrocytic processes. Glia 67, 246–262 (2019).

18. Wang, Y. et al. Accurate quantification of astrocyte and neurotransmitter fluorescence dynamics for single-cell and population-level physiology. Nat Neurosci 22, 1936–1944 (2019).

19. Gähwiler, B. H., Capogna, M., Debanne, D., McKinney, R. A. & Thompson, S. M. Organotypic slice cultures: a technique has come of age. Trends Neurosci. 20, 471– 477 (1997).

20. Gogolla, N., Galimberti, I., DePaola, V. & Caroni, P. Long-term live imaging of neuronal circuits in organotypic hippocampal slice cultures. Nat Protoc 1, 1223–1226 (2006).

21. Bindocci, E. et al. Three-dimensional Ca2+ imaging advances understanding of astrocyte biology. Science 356, (2017).

22. Wang, Y. et al. Automated Functional Analysis of Astrocytes from Chronic Time-Lapse Calcium Imaging Data. Front. Neuroinform. 11, (2017).

23. Arrhenius, S. Quantitative laws in biological chemistry. vol. 1915 (G. Bell, 1915).

24. Schipper, L. A., Hobbs, J. K., Rutledge, S. & Arcus, V. L. Thermodynamic theory explains the temperature optima of soil microbial processes and high Q10 values at low temperatures. Global Change Biology 20, 3578–3586 (2014).

25. Komin, N., Moein, M., Ellisman, M. H. & Skupin, A. Multiscale Modeling Indicates That Temperature Dependent [Ca2. Neural Plasticity https://www.hindawi.com/journals/np/2015/683490/abs/ (2015) doi:10.1155/2015/683490.

26. Clark, J. M. The 3Rs in research: a contemporary approach to replacement, reduction and refinement. British Journal of Nutrition 120, S1–S7 (2018).

27. Bazargani, N. & Attwell, D. Astrocyte calcium signaling: the third wave. Nat Neurosci 19, 182–189 (2016).

28. Thrane, A. S. et al. General anesthesia selectively disrupts astrocyte calcium signaling in the awake mouse cortex. Proceedings of the National Academy of Sciences 109, 18974–18979 (2012).

29. Mori, T. et al. Inducible gene deletion in astroglia and radial glia--a valuable tool for functional and lineage analysis. Glia 54, 21–34 (2006).

30. Paukert, M. et al. Norepinephrine controls astroglial responsiveness to local circuit activity. Neuron 82, 1263–1270 (2014).

31. Wu, Y.-W. et al. Spatiotemporal calcium dynamics in single astrocytes and its modulation by neuronal activity. Cell Calcium 55, 119–129 (2014).

32. Kobe, F. et al. 5-HT7R/G12 Signaling Regulates Neuronal Morphology and Function in an Age-Dependent Manner. J. Neurosci. 32, 2915–2930 (2012).

33. Cupido, A., Cătălin, B., Steffens, H. & Kirchhoff, F. Surgical procedures to study microglial motility in the brain and in the spinal cord by in vivo two-photon laser-scanning microscopy. in (2014). doi:10.1007/978-1-4939-0381-8_2.

34. Guo, Z. V. et al. Procedures for behavioral experiments in head-fixed mice. PLoS ONE 9, e88678 (2014).

35. Kislin, M. et al. Flat-floored air-lifted platform: a new method for combining behavior with microscopy or electrophysiology on awake freely moving rodents. J Vis Exp e51869 (2014) doi:10.3791/51869.

36. Pologruto, T. A., Sabatini, B. L. & Svoboda, K. ScanImage: flexible software for operating laser scanning microscopes. Biomed Eng Online 2, 13 (2003).

37. Maggioni, M., Boracchi, G., Foi, A. & Egiazarian, K. Video Denoising, Deblocking, and Enhancement Through Separable 4-D Nonlocal Spatiotemporal Transforms. IEEE Transactions on Image Processing 21, 3952–3966 (2012).

